# Broad Cognitive and Motivational Enhancement through M_4_-Selective Muscarinic Modulation

**DOI:** 10.64898/2026.11.05.764685

**Authors:** Xuan Wen, Sarah Shehreen, Adam Neumann, Kayla J Temple, Craig W Lindsley, Carrie K Jones, Thilo Womelsdorf

## Abstract

The muscarinic M_4_ subtype of acetylcholine receptors is part of agonist compounds considered treatment options for Schizophrenia and Alzheimers disease, but how M_4_-subtype selective modulation affects cognition and motivation is largely unknown due to dose-limiting side effects, a scarcity of receptor-selective compounds, and missing multi-domain assessments of cognition. Here, we surmount these constraints by utilizing a M_4_-selective positive allosteric modulator (PAM) to assess in nonhuman primates a broad array of endpoints spanning motivational effort control and multiple executive function domains. We found broad dose-dependent performance improvements of set shifting indexing cognitive flexibility, visual search indexing selective attention, antisaccade performance indexing inhibitory control, visuospatial maze learning, and temporal order serial object learning, while working memory span was non-systematically modulated. In addition, the M_4_ PAM reduced effort avoidance in an effort-based choice task documenting a pro-motivational effect. Task-composite performance scores showed pro-cognitive effects across multiple doses and an effective dose range for individual subjects in the absence of adverse dose-limiting cholinergic side effects. Together, the broad pro-cognitive and pro-motivational multi-task profile of M_4_ PAM modulation suggests that the M_4_ receptor is a master regulator of executive cognitive functioning, possibly acting on domain-general neural processes across the prefrontal-striatal circuits.

## Introduction

Muscarinic acetylcholine receptor subtype 4 (M_4_R) has been implicated to mediate pro-cognitive effects in preclinical studies of rodents (Bubser et al., 2012) and nonhuman primates (Lange et al., 2021). In humans, M_4_R’s have been targeted together with the M_1_ receptor by Cobenfy (xanomeline with trospium chloride), a drug FDA approved for reducing positive and negative symptoms in patients with schizophrenia, to improve cognitive functioning (Brannan et al., 2021; McCutcheon et al., 2024; Halassa, 2025; Horan et al., 2025; Sears and Halassa, 2026). While the cognitive effect with M_1_/M_4_ treatment is not evident in the overall population of schizophrenic patients tested with a single dose, cognition is improved among individuals with cognitive deficits at good effect sizes (d ≥ 0.54) (Sauder et al., 2022; Horan et al., 2025; Sears and Halassa, 2026). Remediation of cognitive deficits is consistent with preclinical studies showing M_1_/M_4_ agonists partially recover scopolamine-induced deficits of novel object recognition (Valentino et al., 2026). Together, these findings raise three fundamental questions.

First, what is the specific contribution and scope of the M_4_R subtype in modulating cognition? The dual affinity of xanomeline prevents distinguishing the unique roles of the M_1_ or the M_4_ receptor subtypes for cognition (Javitt, 2025; Leucht et al., 2025). Preclinical evidence suggests that the M_4_ mechanism is particularly prominently implicated in cognitive dysfunctions (McCutcheon et al., 2023) and psychosis (McCutcheon et al., 2024; Saint-Georges et al., 2025) in schizophrenia. Consistent with these findings, cognitive tests in the nonhuman primate (NHP) model show M_4_ -mechanisms enhance inhibitory control (Lange et al., 2021), while M_1_R specific drug compounds increase set shifting abilities but may have negligible attentional effects (Hassani et al., 2023a). What has been missing is an evaluation of the cognitive endpoints of M_4_ selective compounds on a richer set of cognitive functions. Expert consensus in studies of cognitive impairments in schizophrenia (SZ) has proposed seven cognitive domains in the Measurement and Treatment Research to Improve Cognition in Schizophrenia (MATRICS) Consensus Cognitive Battery (MCCB) and its extension Cognitive Neuroscience Treatment Research to Improve Cognition in Schizophrenia (CNTRICS) including separate testing of attention, working memory, inhibitory control, learning and memory (Barch et al., 2009; Nuechterlein et al., 2023). Cognitive performance in most of these domains have unique contributions to the deficits in SZ and bipolar disorder that factor analysis separates into distinct cognitive factors (Schretlen et al., 2013; McCleery et al., 2015; Nuechterlein et al., 2023; Nuechterlein et al., 2025).

Second, does the modulation of the M_4_R subtype enhance cognition and affect motivational functions? The cognitive enhancing effect of Cobenfy in SZ patients with cognitive deficits raises the question of whether targeted potentiation of muscarinic M_4_ receptors can be cognitively beneficial. Classical dopaminergic antipsychotics are not cognitive enhancers and have either small cognitive effects (effect sizes of d<0.23) or impair cognitive and motivational functioning (Feber et al., 2025). This contrasts to apparent improvements reported in subgroups of SZ patients (with d≥0.54 with >1SD lower performance than baseline) (Horan et al., 2025; Sears and Halassa, 2026).

Third, does the positive allosteric modulation of M_4_ receptors allow to discern behavioral effects in a therapeutically meaningful dose range that avoids dose-limiting cholinergic side effects? For a cognitive enhancing drug to have clinical relevance, it should have meaningful therapeutic range. Non-selective cholinesterase inhibition that increases cholinergic tone affects distinct functions at non-overlapping dose levels (Hassani et al., 2023b) and induces gastrointestinal side effects (Ruangritchankul et al., 2021). Similarly, muscarinic agonists often cause dose-limiting adverse effects in patients by acting non-selectively across central and peripheral M_1_, M_2_ and M_3_ receptors (Bodick et al., 1997; Shekhar et al., 2008). Cobenfy mitigates these effects by using peripherally restricted antagonist trospium. An alternative is to utilize positive allosteric modulators (PAMs), which selectively bind at allosteric sites of individual receptor subtypes to potentiate endogenous acetylcholine (ACh) in an activity-dependent manner, minimizing peripheral side effects (Conn et al., 2009).

Here, we address these three questions by assessing cognitive endpoints of VU6008055 (short:VU8055), a highly potent and selective M_4_ positive allosteric modulator (M_4_ PAM), on seven cognitive domains in NHPs. NHPs have similar cholinergic receptor distribution as humans (*see* Suppl. Discussion: *Cholinergic Circuit Differences between Rodent and Primates*), allow for controlled assessment of multiple doses to evaluate the therapeutic dose-range and can routinely perform complex cognitive tasks that relate to ‘real-world functions’ as recommended by FDA for translational clinical impact (Buchanan et al., 2011). We found VU8055 enhanced set shifting, attentional filtering of distractors, inhibitory control, visuospatial (Maze) learning, and temporal order object sequence learning. It also increased the willingness to exert effort for positive outcomes, while having negligible effect on working memory span. This broad cognitive and motivational enhancement extended across two of three doses tested in the absence of adverse side effects.

## Results

The experiment systemically administered (oral) vehicle or VU8055 at 0.3 (low), 1 (medium) and 3 (high) mg/kg doses to six adult male rhesus macaque monkeys before they performed eight distinct tasks assessing executive function domains and motivation. In each session, four of these tasks were performed either from Task Set A or a Task Set B (**Fig. 1A, Fig. S1**). Sessions using *Task Set A* assessed Object Sequence Learning (temporal order working memory), Antisaccade performance (inhibitory control), subjective Effort Control (motivation) and Visual Search (selective attention). Sessions of Task Set B assessed visuospatial Maze learning (spatiotemporal order working memory), Working Memory Span, Probabilistic Feature Learning & Set Shifting and the retention of visuospatial Maze paths by repeating the Maze learning task (**Fig. 1A**). Sessions with Task Set A and B alternated on weekdays ensuring both task sets were used in the same week’s dosing days on Tuesday and Friday. Each subject completed at least seven sessions per dose and task set. Subjects performed the tasks on touchscreen Kiosk Stations in their home cages (Womelsdorf et al., 2021) running the multi-task software platform M-USE (Watson et al., 2023). Subjects initiated the tasks in a fixed task-order and completed all tasks within ∼76-88 min (**Fig. 1A**, **Fig. S1**). Across the 28-week long study, subjects maintained stable weight (**Fig. S2**) and VU8055 did not cause gastrointestinal, somatomotor, or other adverse effects as monitored with a modified Irwin scale (**Fig. S3**,**S4**). Subjects’ performance was assessed with 85 performance metrics obtained across the eight tasks (**Fig. S5**). Vehicle performance was compared to dose conditions using mixed-effects modeling with false discovery rate correction if not stated otherwise.

**Fig. 1.**
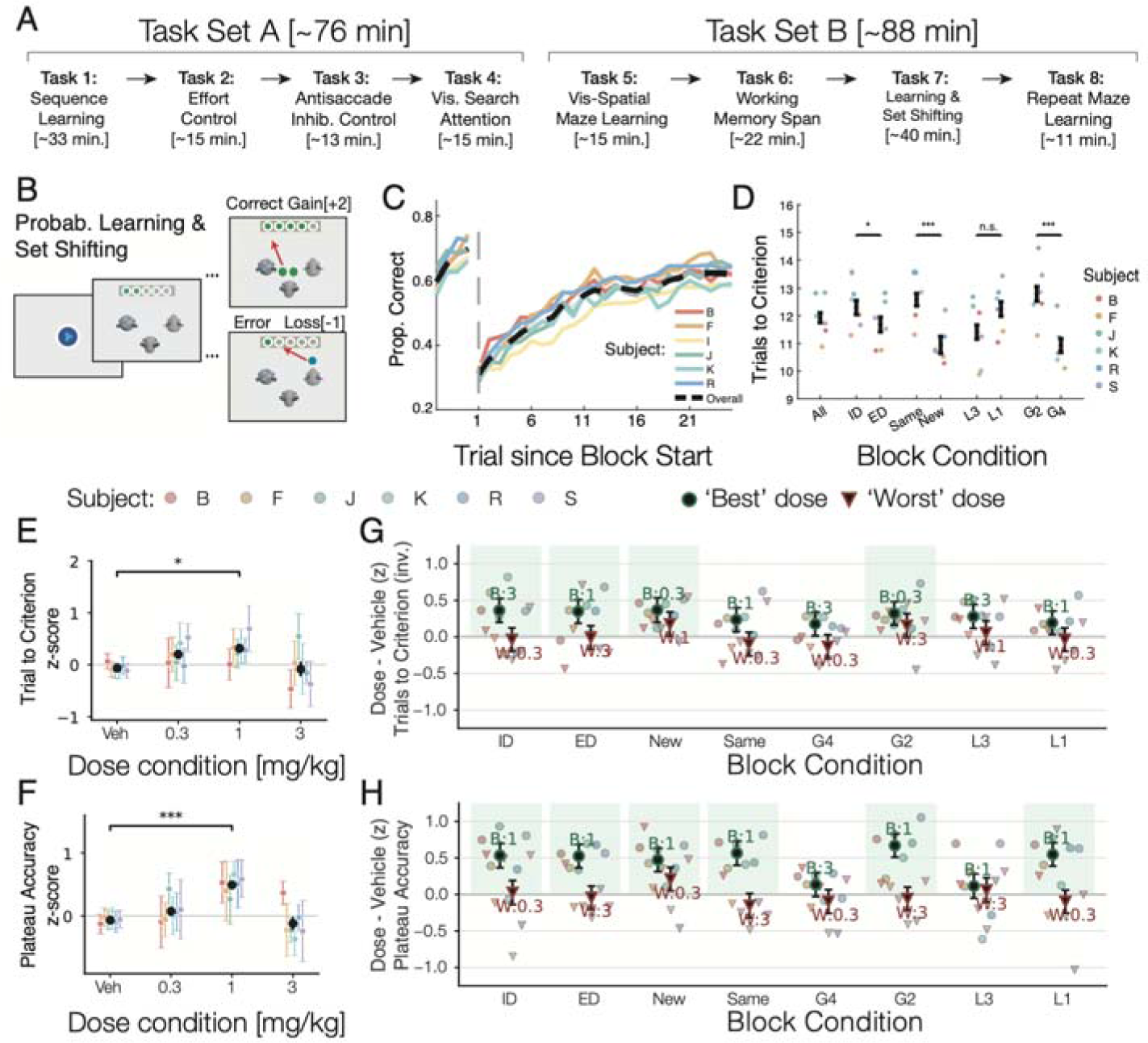
Multi-task assessment paradigm and VU8055 dose-dependent improvement of set shifting performance. **(A)** Subjects performed eight tasks for each dose condition, split into two task sets: Task Set A included Sequence Learning, Antisaccade, Effort Control and Visual Search; Task Set B: included Maze Learning, Working Memory Span, Set Shifting and repetition of Maze Learning. (**B**) The Set Shifting task required subjects choosing one of three objects to receive tokens and avoid losing tokens. The rewarded feature changed every 28-36 trials. (**C**) Subjects learned within 11.96 trials and reached 69.5% plateau accuracy. (**D**) Learning criterion was reached faster when the newly rewarded feature was from a different than the same feature dimension (ED vs ID), when features were novel rather than re-used form the past block (new vs. same), when incentives were higher (4 vs 2 tokens gained for correct responses) (all p<0.05), and there was a tendency to learn faster when errors caused a loss of 3 rather than 1 token (L3 vs L1, p = 0.085). (**E,F**) Vehicle performance norm. learning speed for ED (*E*) (*y-axis*, higher values reflect faster learning) and plateau accuracy (*F*) improved at 0.1 mg/kg. (**G**) Learning speed (higher values reflect faster learning) at the best (circles) and worst (triangles) doses across task conditions. Green shadings denote improvement (p<0.05) at the best dose (session-level LME model). Average (±SE) and individual subjects data are in solid and thin symbols, respectively. (**H**) Same as *G* for plateau accuracy. VU8055 increased plateau accuracy for ID shifts (at 1 mg/kg, beta = 0.56 ± 0.17, p = 0.00073), ED shifts (at 1 mg/kg, beta = 0.55 ± 0.17, p = 0.00085), New target features (at 1 mg/kg, beta = 0.50 ± 0.17, p = 0.0024), with Same features as the previous block (at 1 mg/kg, beta = 0.58 ± 0.17, p = 0.00048), when correct trials gained 2 tokens (at 1 mg/kg, beta = 0.70 ± 0.16, p = 0.000021) and when errors caused loss of 1 token (at 1 mg/kg, beta = 0.58 ± 0.17, p = 0.00049).

### VU8055 improves set shifting and choice accuracy under uncertainty

The Probabilistic Feature Learning and Set Shifting task (Task Set *B,* Task 7) assessed shifting attention sets (**Movie S1**). Subjects learned in blocks of 27-40 trials which object feature is rewarded by choosing objects and using probabilistic token gains and losses as feedback (**Fig. 1B**,**C**). The rewarded feature changed between blocks without an explicit cue. Subjects needed fewer trials to learn (criterion: 75% correct) when a rewarded feature was from a feature dimension that was different to the rewarded feature dimension in the previous block (extra-vs. intra-dimensional shifts: 11.72 vs 12.33 trials, p = 0.029), when visual features of targets and distractors were novel and not used previously as distractors and targets (new vs. same: 10.96 vs 12.62 trials, p = 1.10 x 10^-5), and when the incentives were higher with 4 compared with 2 tokens (Gain-4: 10.92 vs Gain-2: 12.82 trials, p = 5.73 x 10^-5) (**Fig. 1D**; for similar effects on plateau accuracy: **Fig. S6A**). VU8055 improved learning speed and plateau accuracy in blocks with extradimensional shifts (1 mg/kg) (**Fig. 1E-F**) and multiple other set shifting conditions (**Fig. 1G-H**). To discern the net positive effects across performance metrics we extracted the dos with the most consistent improvement (‘best-dose’) and impairment (‘worst-dose’), respectively, across subjects. The set shifting task only provided best-doses and no significant worst-dose effect (green bars indicate best-dose significance), illustrating widespread drug-induced improvements. Subjects learned faster in blocks with intra-dimensional shifts (best-dose 3 mg/kg; beta = 0.38 ± 0.17, p = 0.023) and extra-dimensional shifts (best-dose 1 mg/kg, beta = 0.37 ± 0.17, p = 0.026), in blocks that switched to novel features (best-dose 0.3 mg/kg, beta = 0.38 ± 0.17, p = 0.025), and when blocks offered lower incentives of 2 tokens gained for correct responses (best-dose 0.3 mg/kg, beta = 0.35 ± 0.17, p = 0.037) (**Fig. 1G**). The positive effect of VU8055 on learning speed was paralleled by improved plateau accuracy in the same conditions with best-dose effects consistently at the 1mg/kg dose (**Fig. 1H**). In addition, VU8055 at 1 mg/kg also improved plateau accuracy relative to vehicle in the condition with higher interference when blocks used the same features as the previous block (β = 0.5771 ± 0.165 SE, p < 0.001), and when errors were punished with the loss of 1 token (β = 0.5770 ± 0.165 SE, p < 0.001) (**Fig. 1H**). In finer grained trial-level analysis we confirmed that VU8055 improved at the 1mg/kg dose the ability of subjects to improve performance after error feedback (**Fig S6C**,**D**).

### VU8055 improves attentional filtering of distractors

The Visual Search (Task *Set* A, Task 4) task assessed selective attention with a visual search paradigm that tested how well subjects find a predefined target object in a display with 3-12 distractor objects (**Fig. 2A**, **Movie S2**). Distractors did or did not share visual features with the target i.e. were similar to the target or not. Subjects showed a set-size effect with falling accuracy with increasing number of distractors when they were similar to the target (High feature similarity: beta = −0.0206, p = 1.55 x 10^-55), but not when distractors were distinct from the target, i.e. when the target popped out among distractors (Low feature similarity: beta = −0.0012, p = 0.201)(**Fig. 2B**). VU8055 improved the set-size effect at low (0.3 mg/kg) doses when target-distractor similarity was high, i.e. when search was difficult (beta = 0.372 ± 0.165, p = 0.0239)(**Fig. 2C**), which was confirmed with the best-dose analysis (beta = 0.354 ± 0.157, p = 0.0245)(**Fig. 2D**). There were no or inconsistent drug effects on choice reaction times (**Fig. 2D**, **Fig. S7A**).

**Fig. 2.**
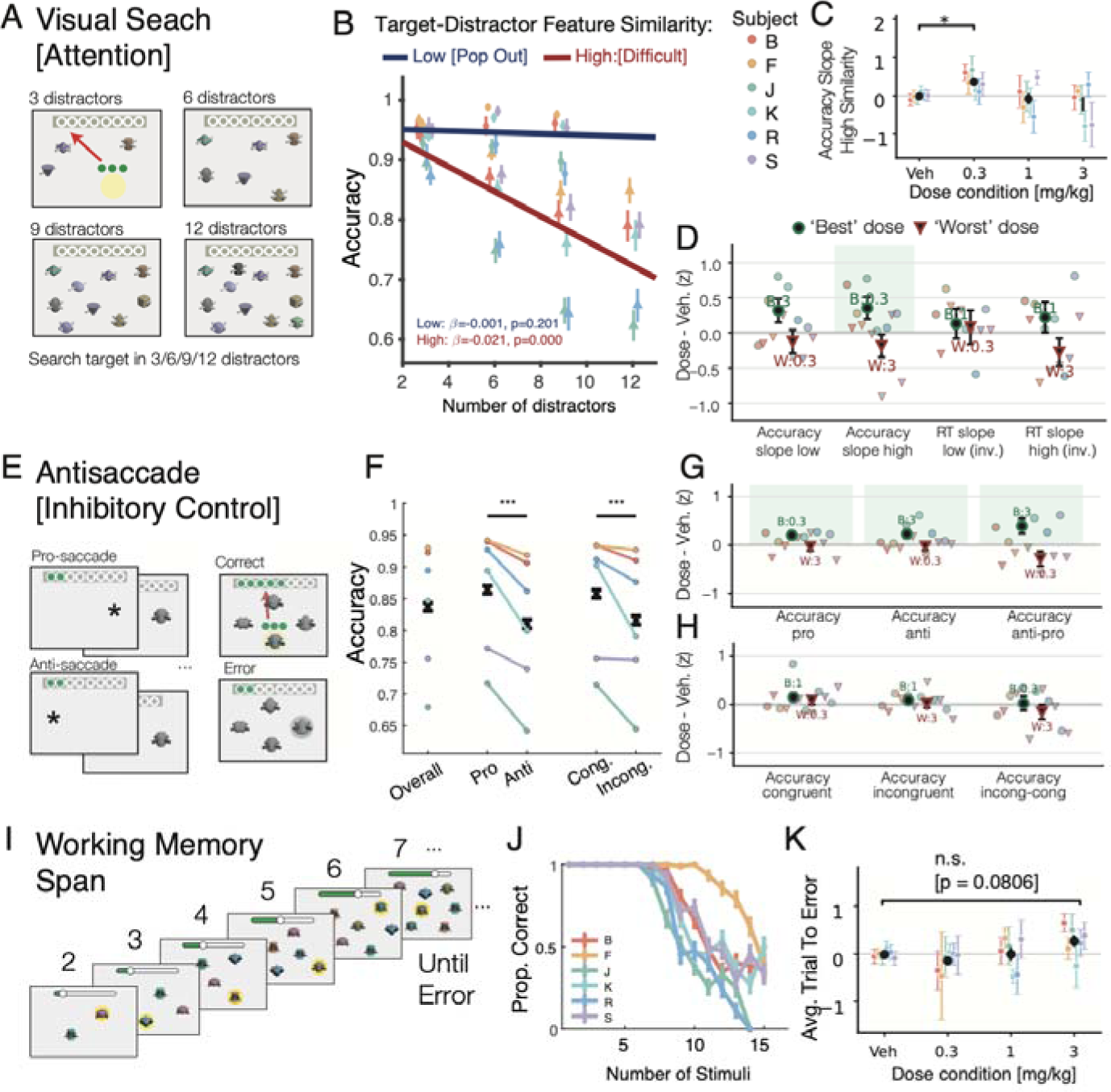
M_4_-selective PAM improved visual search and antisaccade performance but not working memory span. **(A)** Visual Search required control of interference from 3-12 distractors. (**B**) Subjects showed poorer accuracy with more distractors when they were similar to the target (red regression) but not when target-distractor (TD) similarity was low (blue) (LME model, high TD similarity: β = −0.0206, p<0.001; low TD similarity: β = −0.0012, p = .201). (**C**) Vehicle norm. visual search accuracy show improvement at the 0.3 mg/kg dose when TD similarity was high (LME model on accuracy slope: 0.3 mg/kg: β = 0.372 ± 0.165, p = .0239). (**D**) Best (circles) and worst (triangles) doses selectively improved accuracy at high TD similarity (slope: β = 0.354 ± 0.157, p = .0245). (**E**) The Antisaccade task required identifying a target object at the same or opposite side of a flashing cue. (**F**) Accuracy was lower in the anti-than pro-saccade condition (paired t-tests, p<0.001), and when the target was displayed at opposite side in the response display (congruent vs incongruent accuracy: p<0.001). (**G**) The 0.3 mg/kg dose improved accuracy in the pro-condition and 3 mg/kg in the anti-condition (LME: Pro accuracy, 0.3 mg/kg: β = 0.435 ± 0.165, p = .0085. Anti accuracy, 3 mg/kg: β = 0.436 ± 0.166, p = .0088). (**H**) There was no dose effect for the congruency effect. (**I**) The Working Memory Span task required subjects choosing an object not previously chosen in a visual array that added a new object after every choice. (**J**) Subjects committed errors when the display contained ≥6 objects. (**K**) Vehicle norm. trial to an error (indexing working memory span) only showed a trend of improvement at the 3mg/kg dose (LME: average trials to error: 0.3 mg/kg, p = .426; 1 mg/kg, p = .994; 3 mg/kg, p = .081).

### Inhibitory control: VU8055 improves antisaccade performance

The third task (Task Set *A:* Task 3) used an accuracy-based antisaccade task to assess how well subjects inhibited orienting to a peripheral cue that transiently flashed opposite to a target object (anti-saccade condition) compared to a cue at the same location as the target (pro-saccade condition) (**Movie S3**). The target was shown only briefly and followed by a response display with 4 objects (one of them identical to the target) (**Fig. 2E**). Subjects consistently performed poorer for the anti-than pro-saccade condition (anti: 81% vs pro: 86.4%, p < 0.001) (**Fig. 2F**) and also showed a spatial congruency (’Simon’) effect with better performance when the cued target location was congruent with the target position in the response display (incongruent 81.6% vs congruent 85.8%, p < 0.001) (**Fig. 2F**). At 3 mg/kg VU8055 improved anti-saccade accuracy (beta = 0.227 ± 0.087, p = .0095), which exceeded the prosaccade effect (difference anti-prosaccade condition: beta = 0.390 ± 0.156, p = .0125) (**Fig. 2G**). Best dose analysis also showed that VU8055 improved pro-saccade accuracy at 0.3 mg/kg (beta = 0.200 ± 0.083, p = .0160). There was no dose effect on spatial congruency (**Fig. 2H**). Choice reaction times also showed an anti-saccade effect (anti: 1.213 s vs pro: 1.169 s, two-sided paired t-test, t(273) = 9.33, p < 0.001) (**Fig. S7B**).

### No consistent dose effect on working memory span

Task 6 on Task Set *B* assessed working memory (WM) span. Subjects were presented with arrays of objects and had to choose an object in the array that they did not chose previously (**Movie S4**). Within a block of trials, all chosen objects and a novel object were added to the array for subsequent choices, which grew the number of objects subjects held in WM until they committed an error, indexing the limit of their WM span (**Fig. 2I**). Accuracy declined as the number of objects in WM increased, dropping to 75% accuracy after on average 9.33 (SE = 0.81; range: 7.66-13.23) objects, which defines the WM span of subjects (**Fig. 2J**). VU8055 at the high (3 mg/kg) dose showed a tendency to improve performance (p = 0.0806) (**Fig. 2K**) but overall had no systematic effect on WM span accuracy, reaction times, or the WM recency effect (all p > 0.08) (**Fig. S7C-E**).

### VU8055 improved acquisition of the temporal relation of objects

VU8055 improved set shifting, attentional filtering, and inhibitory control, but not WM span. These cognitive functions are required in more complex tasks which we assessed using temporal order object sequence learning (Task Set *A*: Task 1) and visuospatial learning (Task Set *B*: Task 5, 8). First, we tested task how VU8055 affected learning the serial ordering of 4 objects presented together with a distractor (**Fig. 3A**, **Movie S5**). Subjects performed above chance at each ordinal position (**Fig. 3B**) and learned to complete sequences on average within 2.72 ± 0.11 trials. Completion was defined as concluding all 4 objects in the correct order with less than 8 errors in a trial (**Fig. 3C**). After the fifth trial we masked future objects at variable times during the trial to assess prospective working memory for objects at masked ordinal positions (Wen et al., 2026a). Low and medium doses of VU8055 systematically reduced the number of erroneous choices during the first five trials (’early errors’, and the ‘cumulative errors before reaching the learning criterion), and improved accuracy choosing the first object of the sequence across trials (**Fig 3D**,**E**; **Fig S8**). These learning improvements contrasted to the absence of modulation of performance accuracy once learning completed, and to the absence of dose effects on prospective working memory abilities evident in no change of accuracy on masked choices (**Fig 3E**; **Fig S8**).

**Fig. 3.**
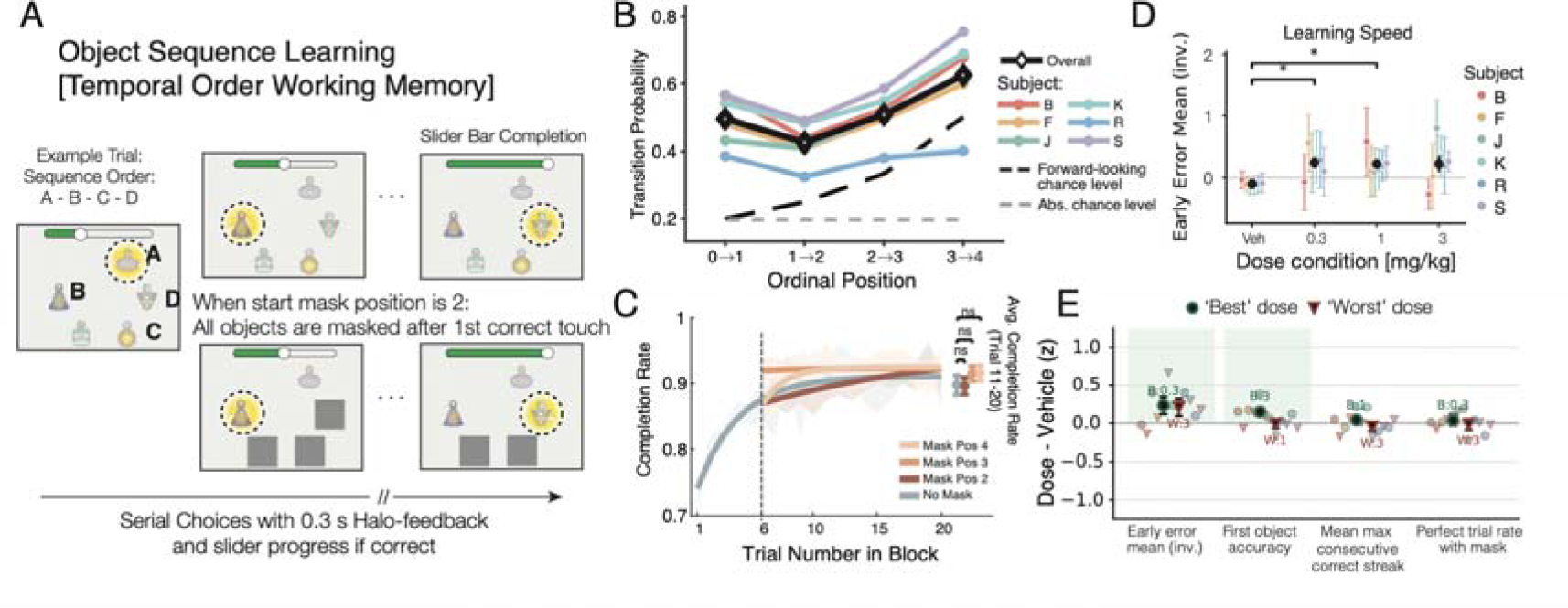
Learning of object sequences improved at low and medium VU8055 doses. **(A)** Subjects learned choosing four objects in a pre-defined A-B-C-D order, ignoring a fifth distractor object. Correct ordinal choices were followed by visual feedback (yellow halo) and progressed a slider. After 6 trials a choice could be followed by a mask of the remaining objects to test for planning. (**B**) Five of six subjects showed above chance accuracy at each ordinal position. On average, transition probabilities were above chance across all ordinal positions (0->1: 0.495; 1->2: 0.425; 2->3: 0.507; 3->4: 0.625; all p < 0.001, LME). (**C**) Sequences were completed at high rates after 6^th^ trial in the no-mask condition and when masking started at position 2, 3, or 4 (No Mask: 0.898 ± 0.015; mask start position 2: 0.896 ± 0.013; position 3: 0.915 ± 0.011; position 4: 0.916 ± 0.012; mask vs no-mask performance was not different, LME, Bonferroni-corrected p >= 0.067). (**D**) Low and medium doses reduced vehicle normalized errors in the first five (no-mask) trials (higher value correspond to less early errors: 0.3 mg/kg: beta = 0.348 ± 0.164, p = 0.0342; 1 mg/kg: beta = 0.326 ± 0.163, p = 0.0448, LME model). (**E**) A significant best-dose effects were found for early learning (reduced errors and improved accuracy of the first ordinal object), but VU8055 did not the proportion of consecutive correct choices or perfectly completed sequences, i.e. without errors). Green shading denotes

### VU8055 improves visuospatial learning and memory

The Maze Learning task (Task Set *B:* Task 5, 8) was an adoption of the Groton Maze learning task used to assess visuospatial learning efficiency and error monitoring in humans (Pietrzak et al., 2008). Subjects were shown a 6x6 grid of tiles and asked to find an invisible hidden path between a start and end tile by following two rules: choose the next tile horizontally or vertically adjacent to the last correct tile, and when committing an error, re-choose the last correct tile (**Fig. 4A**, **Movie S6**). Hidden paths were 14-17 tiles long with 4-6 turns. Correctly chosen tiles were briefly colored green as feedback. Each maze had a unique tile and background color to enable recall of the path when repeating it ∼60 min later in the session in task 8 of Task Set *B*. Subjects learned the hidden path by reducing errors over consecutive attempts and showed memory retention by committing fewer errors in the first trial of encountering a repeated maze at the end of the session (errors per tile in first vs fifth attempt: 1.21 vs 0.68, p = 2.95 x 10^-14; errors in first repeated vs first initial encounter: 0.85 vs 1.21 errors per tile, p = 2.39 x 10^-9) (**Fig. 4B, Fig. S9A-D**). VU8055 at the low dose consistently improved performing the hidden path by (*i*) reducing the rate of perseverative errors (beta = 0.267 ± 0.132, p = 0.0431), (*ii*) by reducing ‘rule-breaking’ errors that violated the task rules (beta = 0.254 ± 0.127, p = 0.0458), and (*iii*) reducing errors committed when the next correct tile lay towards the end-tile rather than away from it (beta = 0.365 ± 0.131, p = 0.0055) (**Fig. 4C-E**). VU0855 also partially improved plateau performance after learning completed as is evident in less errors on the last (5^th^) attempt of a maze at 0.3 mg/kg (**Fig. 4E**). When a maze repeated, the low dose of VU8055 reduced the average error rates of repeat mazes (Repeat mean error: 0.3 mg/kg: beta = 0.30 ± 0.14, p = 0.032), and shortened the average duration needed per tile to complete repeated mazes (Repeat duration per tile: 0.3 mg/kg: beta = 0.31 ± 0.14, p = 0.026) (**Fig. 4F**), but did not benefit and even impaired memory at the 1 mg/kg dose evident by worst dose effect increasing the number of errors during learning of repeat mazes (Repeat first error)(**Fig. S9E**).

**Fig. 4.**
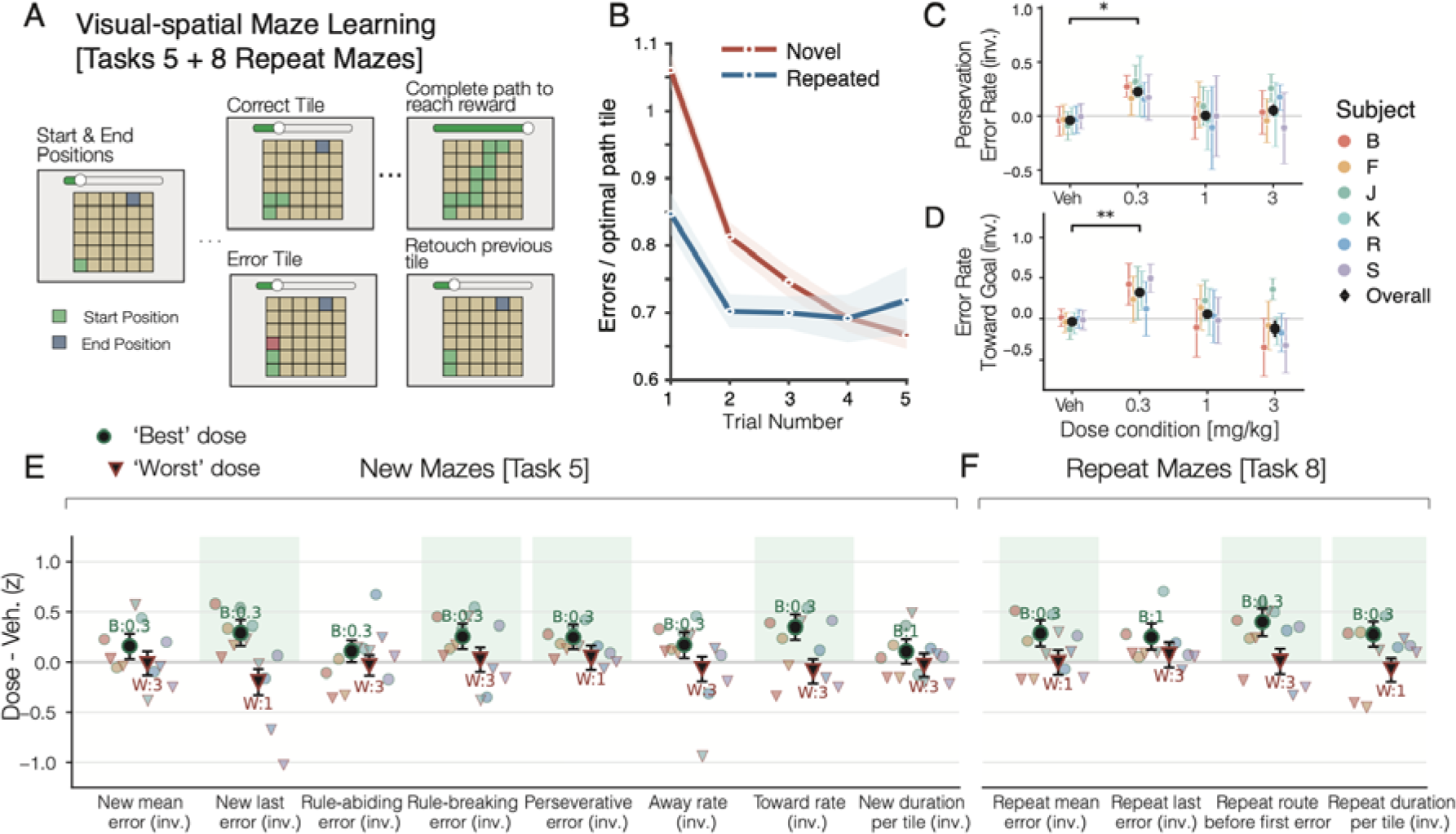
Visuo-spatial Maze learning (Task 5) and retention (Task 8) of hidden Maze paths improved at low VU8055 doses. **(A)** Subjects learned an invisible path from a start to an end tile of a 6x6 grid by choosing adjacent tiles, retouching the last correct tile after an error (correct tiles did not remain visible and are shown for illustration purposes only). (**B**) Learning was reflected in reduced errors over 5 trials (each trial allowing a maximum of 20 errors). Repeating a task-5 maze in task 8 caused less errors indexing memory retention (blue line). (**C**,**D**) The 0.3 mg/kg dose reduced perseverative errors (*C*) and errors when correct tiles laid towards the goal (*D*). Higher values index less errors. (**E**) Best dose effects for task 5 confirm reduced errors at 0.3 mg/kg for different error types. (**F**) Best dose effects for task 8 also show less errors of repeated maze paths. Green shading in

### VU8055 increases willingness to exert effort for reward

We next used an effort-based decision-making task (Task Set *A*: Task 2) assessing motivation. Subjects had to choose one of two virtual balloons and to obtain reward inflate the chosen balloon by touching it as often as indicated by the number of outlines within its confines (**Fig. 5A**, **Movie S7**). The number of touches (outlines) corresponds to physical effort. Effort varied independently of the amount of expected reward that was symbolized by the number of gold-colored tokens shown on-top of each balloon option, causing subjects to weight the possible benefits (tokens) and costs (effort) of choice options. Subjects chose the option with larger token rewards more often when it required less effort, and they chose the option requiring larger effort more likely when it yielded higher token pay off (**Fig. 5B**,**C**). The lower dose of VU8055 reduced effort avoidance, causing subjects to choose a higher effort, higher reward magnitude option more often compared to the vehicle condition (**Fig. 5D**). This effect shows that VU8055 increased the willingness to exert effort for reward, which was evident in a rightward shift of the 50% and the 25% effort-control threshold relative to the vehicle condition by 0.62 and 1.28 additional-effort units, respectively (subject-clustered generalized estimating equation-based threshold tests: p = .031 and p = .010) (**Fig. 5D**,**E**). This finding was confirmed with a linear model that predicted choices by weighting the reward difference and the effort difference of options. The model showed the low dose of VU8055 had no effect on the subjective reward value (0.3 mg/kg: beta = −0.24 ± 0.17, p = .139) (**Fig. 5F**), but reduced effort avoidance (reflected in an increase of the sign-inverted effort avoidance weight; beta = 0.47 ± 0.16, p = .0037) (**Fig. 5G-J**). This positive VU8055 influence on effort control was validated in two additional measures of motivational willingness to choose higher effort options (increases of the reward-effort ratio, and of the motivational breakpoint threshold, **Fig. 5H,I**). These effects were confirmed as best dose effects (Fig. 5J).

**Fig. 5.**
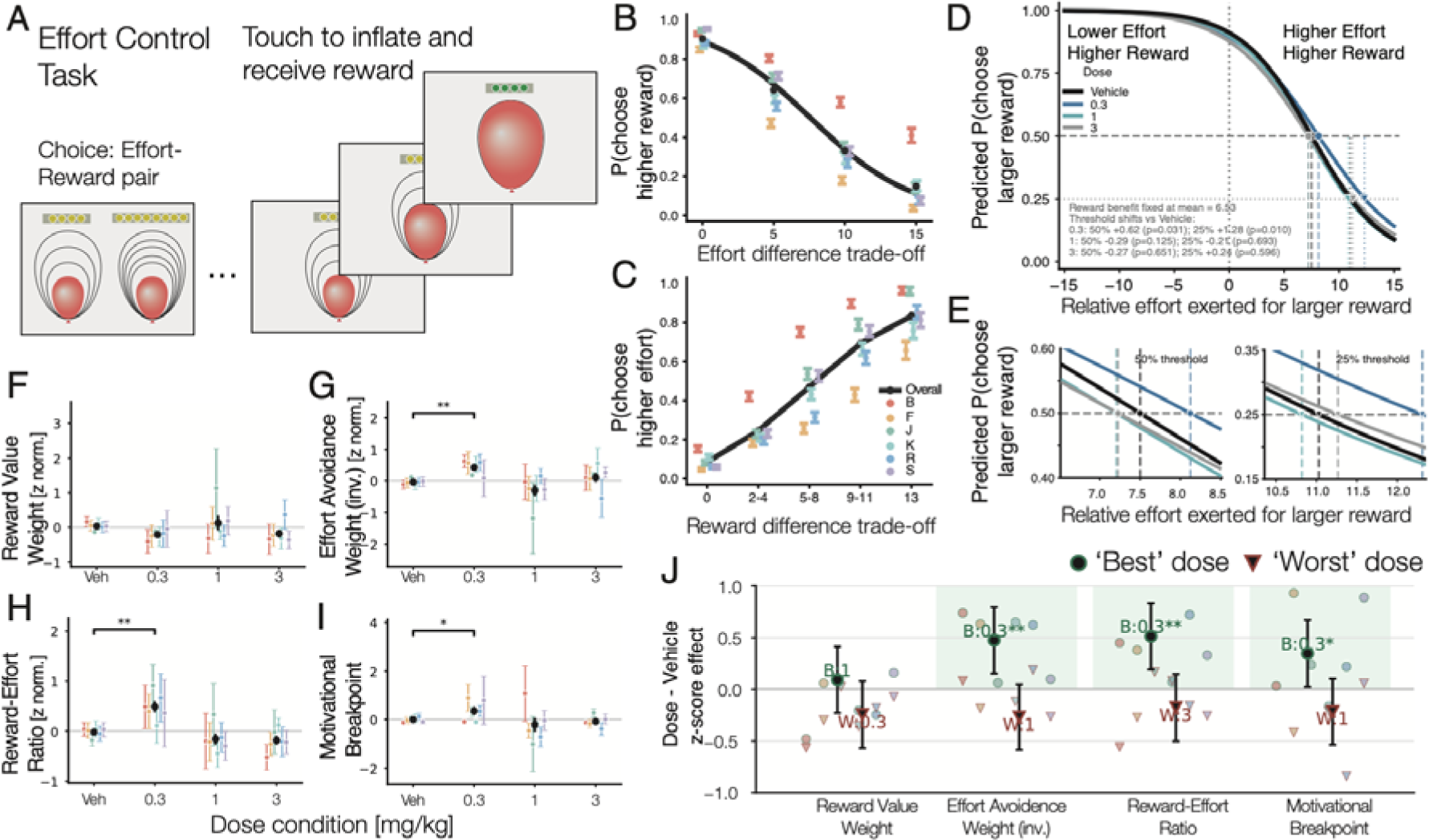
Low dose of M_4_ selective PAM increases willingness to exert effort for obtaining reward. **(A)** The Effort Control task required choosing a balloon with either low or high effort (symbolized by outlines corresponding to number of touches to inflate balloon) and either low or high reward (number of token symbols). Once an effort-reward pair was chosen, touching it virtually inflated the balloon causing the tokens to turn green triggering dispensing fluid reward. (**B**,**C**) Subjects chose the higher reward option (*y-axis*) when it required less effort (*B*) and they chose the higher effort option (*y-axis*) when it paid off larger reward. This pattern shows subjects traded the benefits (rewards) and costs (effort). (**D**) Fitted curves of the average choices of the higher-reward option (*y-axis*) relative to the additional effort that option required. Colors denote dose condition. Dashed grey lines show above 50% of choosing the higher reward option when it required exerting 7.5 more effort (more touch outlines) for obtaining that higher reward. Dotted grey lines denote the 25% threshold. (**E**) Subsection of D zoomed in on the 50% and 25% threshold visualizing that 0.3 mg/kg VU8055 increased the amount of effort subjects exerted for obtaining higher rewards. (**F)** Best doses (circles) and worst doses (triangles) show that 0.3 mg/kg made the effort coefficient less negative, indicating reduced effort avoidance, and increased the 25% cost-reward effort threshold, indicating less avoidance, i.e. a greater willingness to choose a higher-effort option for additional reward. Green shading denotes p<0.05 (session-level LME).

### No influence of baseline performance on the strength of dose effects

We next tested whether drug-induced cognitive and motivational improvements were more pronounced in lower performing subjects. However, we found that baseline performance levels of the six subjects of our sample did not show correlations with drug-induced improvements in any of the tasks (all n.s., FDR corrected, **Fig. S10**). There was only one metric, the response time slope in the visual search task in the low target-distractor similarity condition, for which faster baseline response times was associated with stronger dose-induced response speed increase (β = 1.017, 95% bootstrap CI [0.536, 1.930], p = .0496, FDR corrected).

### VU8055 has a net-positive task composite effect and subject-specific dose efficacy

To summarize the main findings, we extracted the best- and worst-dose effects of the core performance metrics of each task category. This showed best-dose improvements in six of the seven task categories, worst-dose impairments in two task categories, and no modulation of WM span and subjective reward value in the effort control task (**Fig 6A**). The effect sizes ranged from ∼+0.5 z-normalized standard deviations (z-std) for effort control and set shifting plateau performance, to ∼+0.3 z-std for attentional filtering and inhibitory control antisaccade performance, and ∼+0.25 z-std for set-shifting learning speed, visual-spatial maze learning, and object sequence learning. For these effects the best dose included to varying degrees the 0.3 and the 1 mg/kg dose and for a few metrics also the 3 mg/kg dose. To go beyond best-/worst-doses, we quantified the breadth of dose effects for each task by calculating the average effect per dose as a *task-composite dose effect* using the 37 accuracy-based performance metrics that showed a statistical effect at any of the three doses (**Fig. 6B**). The *task composite effect* was net-positive at 0.3 and 1 mg/kg, but not at 3 mg/kg (two-sided one-sample t-test of six subject-level equal-task means; 37 metrics; 0.3 / 1 / 3 mg/kg: p = 0.002 / 0.005 / 0.346). A similar result was evident including in the analysis all 67 accuracy-based performance metrics irrespective of whether they showed individual dose effects (**Fig. S11A**). Qualitative inspection showed that the task-specific dose effect scores of the maze learning, sequence learning and effort control peaked at the low dose, while the composite scores of the visual search and set shifting task peaked at the medium dose and the inhibitory control metrics of the antisaccade task peaked at the high dose.

**Fig. 6.**
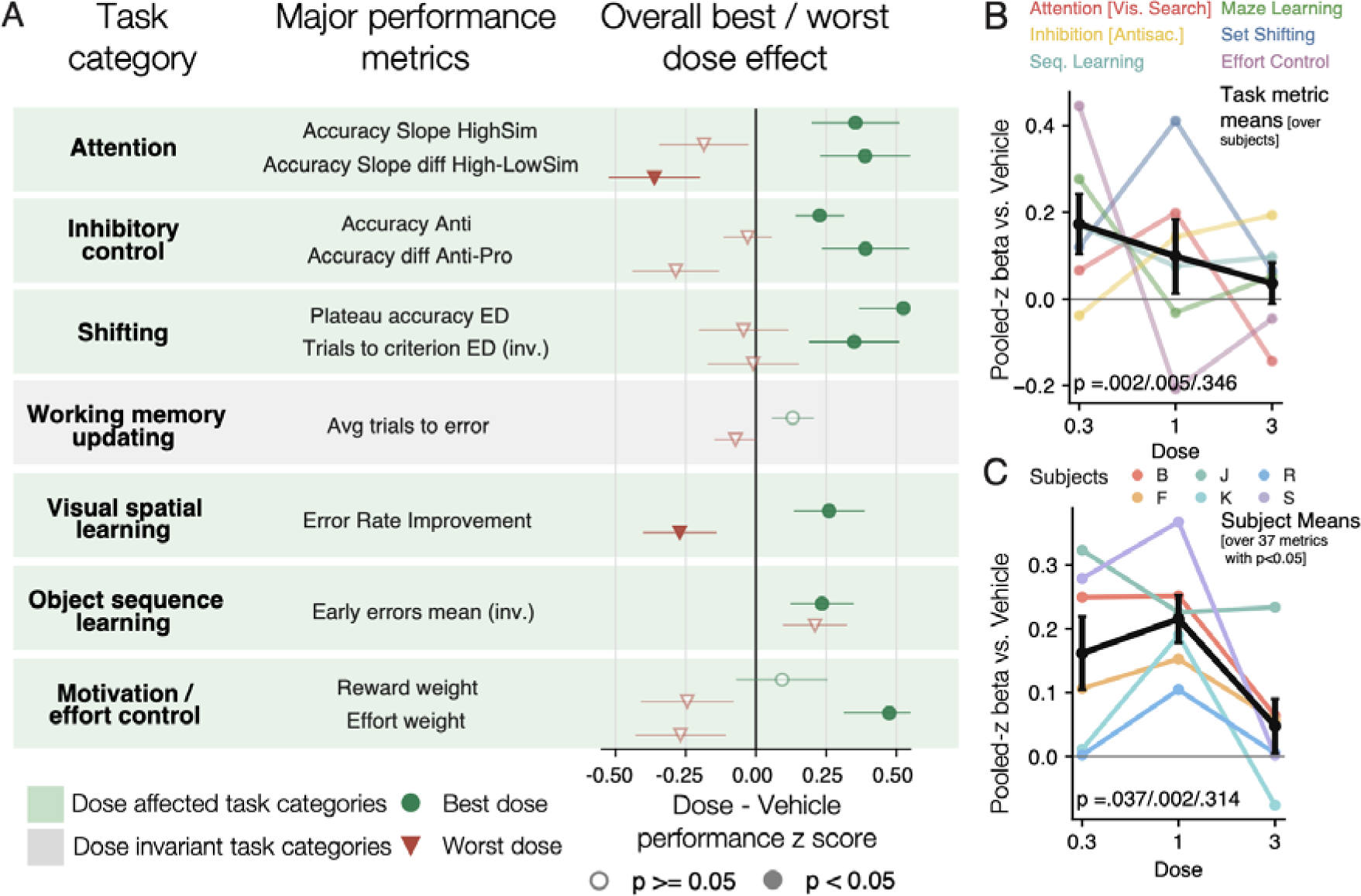
VU8055 improved performance in six of seven task categories. (**A**) The average best and worst dose vs. vehicle effects (green circles and red triangles, respectively) for the core performance metrics (*middle* column) of each task (*left* column). (**B**) Task-composite dose-response profile. Colored lines show task-specific mean pooled-z mixed-model coefficients for each dose relative to vehicle (*x-axis*), averaged across all accuracy-based metrics of each task. Black line is the mean across tasks; error bars are SEM across tasks. Values at each dose were tested against zero using two-sided one-sample t-tests (0.3/1/3 mg/kg: p = 0.002/0.005/0.346). (**C**) Subject specific average dose effects (in color) and the average subject-level dose effect of VU8055 (black line). The subject-specific effect is the mean pooled-z value at a given dose minus the mean of that subject’s vehicle. The black line is the mean across subjects; error bars indicate SEM. Values at each dose were tested against zero using two-sided one-sample t-tests (n = 6 subjects; 0.3/1/3 mg/kg: p = 0.037/0.003/0.314).

Next, we calculated subject-specific dose efficacy profiles across task metrics, using the same 37 metrics as for the main task-composite analysis but averaging the results across metrics and tasks. First, we found that on average subjects showed across metrics a net-positive cognitive effect at 0.3 and 1 mg/kg, but not at 3 mg/kg (two-sided one-sample t-test; n = 6 subjects; 0.3 / 1 / 3 mg/kg: p = 0.037 / 0.002 / 0.314) (**Fig. 6C**; see **Fig. S11B** for similar results using all 67 metrics). Second, qualitative inspection of the results suggests that some subjects showed beneficial net-positive effects at low and medium dose (Subjects B, S, and F) or at all three doses (subject J), while two subjects showed net-benefits across metrics exclusively at the medium dose (subjects K and R) (**Fig. 6C**). To understand the magnitude of subject-specific effect sizes in our sample, we extracted for each subject the strongest positive dose effect and then averaged these effects across metrics and subjects. This descriptive approach resulted an average subject-specific best-dose effect +0.33 z-std across all 85 metrics (**Fig. S12**).

### Days of testing has negligible effects on dose effects

Task performance with dose administration was conducted on Tuesdays and Fridays with dedicated weeks with vehicle administration on Tuesdays and Fridays similar to a previous study (Lange et al., 2021). This design allowed addressing two questions to test for the generalizability of the results. First, we tested whether dose effects on performance metrics were similar on Tuesdays and Fridays and found overall similar effects with no statistical differences (**Suppl. Results** ‘*Similar dose effects on Tuesdays and Fridays’,* **Fig. S13**). Second, we tested whether performance effects with vehicle administration on Tuesdays/Fridays were similar to performance metrics with vehicle administration on Monday/Wednesdays/Thursdays and found overall similar effects with no differences between the vehicle day categories (**Suppl. Results** ‘*Similar performance with vehicle administration on Tuesdays/Fridays and on Mondays/Wednesdays/Thursdays’,* **Fig. S14**,**S15**).

## Discussion

We found that VU8055 improved executive cognitive functions and effort control across a broad range of domains assessed across six of seven tasks (**Fig 6A**), a net-positive task composite scores at the low and medium dose (**Fig 6B**) and an average net-positive pro-cognitive effect for individual subjects (**Fig 6C**).VU8055 improved (*i*) learning and accuracy during set-shifting, (*ii*) visual search when the interference from distractors were high, (*iii*) inhibitory control in antisaccade over pro saccade trials, (*iv*) learning of temporal order object sequences, (*v*) visuospatial maze learning by reducing persevere and rule-breaking errors, (vi) improved visuospatial recognition memory as inferred from faster and less erroneous performance of repeated mazes at the end of the session, and (vii) it reduced the tendency to avoid higher effort options. The metric not systemically improved by VU8055 was working memory span which showed a trend for improvement at a higher dose that was outside of the net-positive task-composite effect. These findings emerged in the absence of negative adverse gastrointestinal side effects.

### Origins for broad ‘domain-general’ cognitive motivational improvements

The breadth of the cognitive enhancing effect of a receptor specific compound in NHP’s is unprecedented and suggests that the M_4_ receptor mechanism serves as a master regulator within a network supporting multiple executive functions. The cognitive effects with the M_4_ PAM can be summarized as improving cognitive flexibility and interference control of goal representations. Improved cognitive flexibility was evident from greater ability to learn and update top-down goal representations during feature-reward learning and set shifting (**Fig. 1**), object sequence learning (**Fig 3D,E**), and visuospatial learning (**Fig. 4C-E**). Improved interference control of goal representations was evident in better filtering of distractions during visual search (**Fig. 2C,D**) and stronger attentional control of attentional capture by peripheral stimuli in the antisaccade task (**Fig. 2G**). The brain network supporting cognitive flexibility, interference control, and also motivational control will entail the prefrontal-striatal-thalamic network whose core function can be described as the adaptive selective filtering of goal-relevant information (reviewed in (Pennartz et al., 2009; Pennartz et al., 2011; Cools and Arnsten, 2022; Pathak et al., 2025)). M_4_R’s are expressed at low levels in lateral and medial prefrontal cortex of NHPs where they exert a net-inhibitory influence on pyramidal cells and regulate D_1_-receptor mediated activity (*summarized in* **Fig. S16**) (Tsolias et al., 2026). More prominently expressed are M_4_R’s at fronto-strital synapses and within the striatum in NHP as shown with autoradiography (Li et al., 2022), with rodent studies revealing M_4_R expression on cholinergic interneurons and pre-synaptically on D1-expressing medium spiny neurons (*summarized in* **Fig. S17)** (Brann et al., 1988; Buckley et al., 1988). These distributions of the M_4_ receptor are consistent with a broad effect on regulating fronto-striatal network functions, spanning reinforcement learning and set shifting (Treuting et al., 2025), selective attention (Arcizet and Krauzlis, 2018; Banaie Boroujeni et al., 2020), inhibitory control during antisaccade performance (Koval et al., 2011), sequence learning (Tian et al., 2024), visuospatial learning (Berke et al., 2009; Woolley et al., 2013; Varin and de Kerchove d’Exaerde, 2025), and effort control (Shenhav et al., 2017; Salamone and Correa, 2024). The notable exception to the positive effect of VU8055 on performance in these domains was the WM span measure, which was on average not modulated, although the statistical trend pointed towards a relative improvement (**Fig. 2K**). This finding is consistent with the literature that has more directly implicated the M_1_ and M_3_ receptors in working memory modulation (*see* **Suppl. Discussion:** *Working memory modulation with M_1_ and M_1_/M_3_ receptor specific compounds and nonselective muscarinic compounds*). Updating the content of a growing working memory will depend on recurrent NMDA-mediated prefrontal cortex mechanisms where M_4_R’s activity has been shown in NHP to induce a net-increase of inhibition relative to excitation (Tsolias et al., 2026). Such a reduction of frontal cortex excitation/inhibition ratio could support stabilizing top-down representations by filtering activity coding for distracting inputs but may have limits in supporting the simultaneous maintenance of multiple WM items that is needed in the WM span task. In summary, the broad cognitive and motivational improvement we observed suggest that the M_4_ receptor serves as a master regulator that acts on fronto-striatal information processing during goal-directed behavior.

### Translational significance to clinical conditions in humans

Our study introduces a new multi-task assessment approach that adopted tasks that are similarly used in human subjects with neuropsychiatric disorders. This cross-species connection of the tasks suggests that the cognitive and motivational improvements seen in healthy NHPs have potential relevance for treatment development in humans.

First, attention, set shifting and inhibitory control are executive function domains have broad transdiagnostic relevance, because they are dysfunctional not only in schizophrenia but also in MDD, AD, and other disorder (Gualtieri and Morgan, 2008; Millan et al., 2012; Schretlen et al., 2013; Shi et al., 2021). For SZ and MDD, psychopathological changes can be found across the striato-thalamo-cortical network that supports multiple cognitive domains (Leucht et al., 2025) with consistent evidence for reduced muscarinic M_1_/M_4_ expression in prefrontal cortex, anterior cingulate cortex, striatum and hippocampus (Zavitsanou et al., 2004; Scarr et al., 2007; Saint-Georges et al., 2025). In addition, the fronto-striatal-thalamic network not only supports executive cognitive functions but is closely related to psychosis across psychiatric disorders with positive symptoms (Braun et al., 2026).

Second, we found the M_4_ PAM improved effort control in a task that was adopted from human studies reporting that subjects that showed higher effort control more likely benefitted from treatment in schizophrenia (Gold et al., 2013) and major depression (Berwian et al., 2020). A lack of drive (avolution) and motivational apathy are clinically relevant symptoms in patients with SZ and MDD (Valton et al., 2025; Periche-Tomas et al., 2026; Pillny et al., 2026), with dopamine-antagonizing antipsychotic medications exacerbating a low-effort biases of choices (Salamone and Correa, 2024). These motivational deficits have been found in subgroups of patients to correlate with other negative symptoms including anhedonia and asociality, which emphasizes the potential translational clinical relevance of our findings (Kirkpatrick et al., 2006; Gold et al., 2013; Barch et al., 2014; Cooper et al., 2019).

Third, we found the M_4_ PAM improved performance in complex compound tasks such as Object Sequence Learning the Visuospatial Maze Learning. Complex compound tasks depend on the coordination of multiple cognitive subcomponent processes as they are common in everyday tasks in the real world. Successful object sequence learning depend on selective attention, working memory of recent choices, the use of a strategy following of task rules (e.g. do not choose diagonal tiles in the maze task, and rechose the last correct object after an erroneous choice in the object seq. learning task) and learning of the temporal (sequential) structure of objects (sequence learning) and locations (maze task). The coordination of these multiple subcomponent processes goes beyond narrow experimental tasks and suggests that the M_4_R selective PAM performance enhancement of these tasks may translate to real-world cognitive function improvement. Such a relevance for real-world functioning has been formulated as one FDA guideline for evaluating treatment options for cognitive impairments in human clinical trials (Buchanan et al., 2011).

### Multi-domain assessment of cognitive endpoints

The results of this study were possible by administering eight tasks to rhesus monkeys, alternating sessions with 4 different tasks per day. This multi-task design was conceived to match the breadth of assessments in digital non-verbal batteries for humans such as the CogState Computerized Battery that is designed to match the MCCB MATRICS battery (Pietrzak et al., 2009; Nuechterlein et al., 2025). Such multi-task assessment of multiple domains is pivotal to judge the breadth of pharmacological effects. Recent meta-surveys of non-invasive brain stimulation treatments (Begemann et al., 2020; Grover et al., 2023) and cognitive remediation training (Fitapelli and Lindenmayer, 2022) report of robust effects in specific cognitive domains that suggests a selective impact on cognition that would not be discernible without broad cognitive testing. In NHPs assessments of 2 or more tasks has been rare; we surveyed 19 available NHP studies testing muscarinic compounds on cognition and found that they used either two or less tasks to assess two cognitive domains at most (*summarized in* **Fig. S18**). Together, these studies so far have suggested that M_4_ and M_1_/M_4_ compounds recover scopolamine impaired inhibitory control, learning/set shifting and working memory performance (**Fig. S18**) (Vardigan et al., 2015; Lange et al., 2021; Galbo-Thomma et al., 2024). The multi-task approach we introduced in this study allowed in a single study to elucidate the breadth of the effect, showing enhancements across multiple cognitive domains through multiple task metrics per domain (red arrows in **Fig. S18**), and revealed a broad modulatory effect of the M_4_ receptor that previous single-task studies couldn’t establish.

### Conclusion

Recently, the co-administration of xanomeline (agonist for M_1_+M_4_Rs) and trospium of inpatients with SZ has been reported to cause a rapid response that is qualitatively described as ‘cognitive awakening phenotype’ in half of patients with prominent negative symptoms (Sears and Halassa, 2026). Patients showing this effect appear more engaged with their environment, motivated to address their needs, and structured in their thinking. While this qualitative description is limited and awaits further quantification, it bears resemblance to the broad cognitive and motivational improvements triggered by M_4_ selective modulations in healthy NHPs. In summary, our study provides direct evidence that the M_4_ receptor mechanism is a key mediator of adaptive cognitive and motivational functioning in primate brains.

## Materials and Methods

### Ethics statement

All animal and experimental procedures complied with the National Institutes of Health Guide for the Care and Use of Laboratory Animals and the Society for Neuroscience Guidelines and Policies and were approved by the Vanderbilt University Institutional Animal Care and Use Committee (IACUC) under approval number M1700198.

### Subjects and apparatus

Six male rhesus macaque monkeys were used in this study, 11-15 years of age, and weighing 11.3-11.7 kg (Subject B: 11 yrs/11.3 kg; Subject F: 13 yrs/11.8 kg; Subject J: 14 yrs/11.3 kg; Subject K: 13 yrs/11.4 kg; Subject R: 15 yrs/11.6 kg; Subject S: 11 yrs/11.7 kg). Some of the subjects were part of previous studies with cholinergic modulation (Hassani et al., 2023b; Hassani et al., 2023a) but none of the subjects received drugs for >12 month prior to this study. Subjects performed the experimental tasks in their housing cages using cage-mounted touchscreen Kiosk stations (Womelsdorf et al., 2021). Visual display, behavioral response registration, and reward delivery were controlled by the Multi-Task Suite for Experiments (M-USE), an open-source Unity^®^-based software platform integrated with a touchscreen, a video monitoring system, and reward delivery hardware (Watson et al., 2023). The touchscreen was an Elo 2094L 19.5″ LCD with a refresh rate of 60 Hz and a resolution of 1,920 × 1,080 pixels. Subjects were on a water control schedule and received on average 20-25 ml/kg fluid as reinforcement during task performance on weekdays and as supplement on weekends.

### Multi-task assessment paradigm

For the duration of this study, each subject performed four tasks in each session on weekdays on the touchscreen Kiosk stations mounted to their home cage. The four tasks were either from *Task Set A* or *B* (**Fig 1A**). In one week, *Task Set A* ran Monday, Wednesday and Friday and *Task Set B* ran Tuesday and Thursday. In the subsequent week *Task Set B* ran Monday, Wednesday and Friday and *Task Set A* ran Tuesday and Thursday. This biweekly pattern alternated throughout the 28 weeks of this study and ensured that Tuesdays and on Fridays within a week used not the same task set but both, Task Set A and B. Tuesdays and on Fridays were dose-testing days and in each week the same dose condition was used. Thus, within a week, the same dose was tested on all eight tasks (*see* below).

Task Set’s A and B consisted of 4 separate tasks that subjects performed within the same testing session in on average 76 minutes (Task Set A) and 81 minutes (Task Set B) (**Fig. 1A**). A session started by presenting the subject four thumbnail images that corresponded to the individual tasks as shown in **Fig. S1**. Subjects touched a task-thumbnail to initiate a task. At the beginning of a session only the first of the thumbnail images was not greyed out, so that subjects had to start the first task by touching its thumbnail. After completing the first task the task-selection screen showed all four task thumbnails, greying out all thumbnail images but the image for the next task (**Fig. S1**). The task order was fixed throughout the experiment with all subjects following the same task order in order avoid variability from subjective preferences for tasks (which was not assessed).

In Task Set A, the Task 1 was an Object Sequence Learning task with performance lasting ∼33 min. Five different 4-object sequences were performed, each with a unique set of objects. Every object set was presented 20 times, each time allowing the subjects to complete the sequence with maximally 8 errors. Task 2 was an Effort Control task with an average duration of 15 min. The task presented for 80 trials two options associated with different degrees of reward outcomes and different degrees of effort required to obtain those rewards. Once subjects chose a reward-effort option, a virtual balloon needed to be inflated by touching its outlines to cashout the reward. Task 3 was an accuracy-based Antisaccade task and was performed for ∼13 min. The task presented in 100 trials visual targets at peripheral locations that were either at the same (pro-) or different (anti-) side of a briefly flashing cue. Task 4 was a Visual Search task. Performance lasted ∼12-15 min. during which 150 trials were presented where a target was shown among 3-12 distractor objects which were either different or similar to the target (i.e. either shared or did not share features with the target).

Task Set B consisted also of four separate tasks (tasks 5-8) with task 8 repeating task 5 to test memory retention (**Fig 1A**). Task 5 was a visuospatial Maze Learning Task performed for ∼15 minutes. Subjects learned four different mazes. Each maze was presented for five trials. A maze path spanned 14-17 tiles from start to end with 4-6 turns within a 6x6 grid of tiles. Each maze of task 5 had a unique tile coloring and background image to allow testing their retention later in task 8. Task 6 was the Working Memory Span task and was performed for ∼20-27 minutes, involving 25 separate blocks. A block ended when subjects committed an error at their working span limit. Task 7 was a probabilistic Learning and Set Shifting Task lasting for ∼38-45 min. In each session 17 set shifting blocks were presented, each with 27-40 trials before a block switched and the rewarded feature changed. Task 8 tested the retention of two maze paths of task 5 plus the learning of two new mazes. Performance lasted ∼11 min. The ordering of the repeated mazes differed from the order of the mazes in task 5.

The duration of the individual tasks was chosen for three reasons: (*i*) to ensure that subjects completed all four task of a set consistently on every day of the week, which showed consistent cognitive and motivational engagement and ensured similar fluid intake from positive reinforcement across weekdays; (*ii*) to ensure similar durations for performance of Task Set A and B; and (*iii*) to collect sufficient number of trials and blocks for robust assessment of performance metrics as inferred during training and from previous experience (Wen et al., 2026b). The study began after subjects were trained on the individual tasks and showed stable above-chance behavior. One NHP (subject R) was not able to perform the Maze Learning task (task 5 and 8). For this subject a separate Task Set B was used to ensure similar timing of the onset of task 6 and 7.

Five tasks of the multi-task assessment paradigm used the same type of multidimensional, 3D rendered Quaddle objects (Wen et al., 2025): Object Sequence Learning (Task 1), Antisaccade (Task 3), Visual Search (Task 4), Working Memory Span (Task 6) and the Probabilistic Learning and Set Shifting Task (Task 7). Quaddle objects can vary in up to ten feature dimensions (e.g., shape, color, body pattern, different arm orientations, the presence of a head, etc.), each with more than ten possible feature values. For the tasks using Quaddle objects, novel sets of objects were generated for each experimental session, if not otherwise stated, to avoid effects of previous exposure to object feature constellations on learning and working memory performance.

Each task was associated with a task-specific thema for textured background images. Background context images were generated with DALL·E 2 (OpenAI) using text prompts for obtaining images from categories that were fixed for a given task. For example, the Object Sequence Learning task used rocks as background image category (*see* **Movies S1-S7**). Color filters were applied to images to obtain a larger number of distinct context backgrounds, ensuring each color filter maintained a fixed luminance value of 50. The colors were selected by evenly spacing them around the CIELAB color wheel, ensuring a diverse range of hues. The brightness of the background images was adjusted to be ≤50% of the HSL scale.

### Pharmacological design and dose conditions

VU6008055 (in short: VU8055) was synthesized in house at the Warren Neuroscience Drug Discovery Center. VU8055 is a highly potent and efficacious positive allosteric modulator (PAM) for the M_4_ receptor that binds to an allosteric site, i.e. site spatially distinct and less preserved than orthosteric site across subreceptors, and potentiates the effects of ACh activity (Engers et al., 2025). VU8055 is the most potent and efficacious of the VU M_4_-selective PAMs (VU8055: EC_50_ = 63.5 nM, 89% efficacy; VU7485: EC_50_ = 102 nM, 74% efficacy; VU7154: EC_50_ = 1000 nM, 57% efficacy) (Bubser et al., 2014; Wood et al., 2017; Schubert et al., 2019; Engers et al., 2025).

VU8055 was tested in a within-subject dose design with orally administered doses of 0 mg/kg (vehicle), 0.3 mg/kg, 1 mg/kg, and 3 mg/kg. Vehicle consisted of 18 g of yogurt and 2 g of honey mixed in a paper cup for oral administration. On compound delivery days, VU8055 was mixed in with the yogurt and honey. Vehicle or vehicle plus compound were given via oral administration 30 minutes prior to the start of testing. Subjects waited in front of the touchscreen and consistently started performance once the task selection screen was presented (**Fig S1**), ensuing similar timing of dose administration and performance of each task. Three of the NHP’s were tested at 9:00 am +/- 5 minutes and three NHP’s were tested at 11:00 am +/- 5 minutes.

NHPs were given the Vehicle on Monday, Wednesday, and Thursday, and on Tuesday and Friday they were given Vehicle combined with 0, 0.3, 1 or 3mg/kg of compound VU8055. In a given week, the Tuesday and Friday dose condition was identical, i.e. each of the four dose conditions was repeated seven times over the 28-weeks duration of the study. This dosing regimen ensured that each week the same dose condition was used in one session with Task Set A and in Task B, which differed on Tuesdays and Fridays. To acclimate and test for dosing effects, week 1, 2, 3 and 4 consisted of 0, 0.3, 1 and 3 mg/kg concentration dosing events respectively and the next 24 weeks were pseudo randomly distributed across the remaining weeks of the study (**Fig. S2A**). A total of two dosing events repeated seven times for each concentration and six weeks of vehicle only were conducted for the 28-week study.

The six NHPs were weighed every Monday, and the volume of compound was calculated for each weeks designated concentration. Body weights were stable across the 28 weeks with changes of on average −0.01 kg (−0.1%) between the first and final dosing session, with an exception of two subjects showing drops of weight in the last two weeks of the study due to medical reasons unrelated to the study (**Fig. S2B**).

## Monitoring of adverse side effects

Subjects were monitored during task performance with a video monitoring system mounted within the Kiosk touchscreen stations and streamed to a web browser on a tablet (*see* also **Movies S1-7**). Immediately prior to the start of the task performance and once all tasks were completed, subjects’ status, behavior, and appearance were rated by an experienced research specialist on a formal modified Irwin test that scored possible cholinergic effects on the autonomic and somatomotor systems (**Fig S4**,**5**). Throughout the 28 weeks of dose testing there was not a single instance when ratings were not 0 (no change), i.e. there were no slight (rating 1) or significant changes (rating 2) noted. The absence of side effects contrasts with slight changes and early-dose effects in previous studies using other drug compounds (Hassani et al., 2023b; Hassani et al., 2023a).

### Task paradigms and task-specific performance metrics

The multitask assessment paradigm comprised eight tasks that provided 85 task-specific metrics - listed in **Fig. S5** - to score performance (8 + 4 + 12 + 8 + 13 + 7 + 28 + 5 metrics across Tasks 1–8, respectively): 8 metrics for Task 1, Object Sequence Learning; 4 for Task 2, Effort Control; 12 for Task 3, Antisaccade; 8 for Task 4, Visual Search; 13 for Task 5, novel visuospatial Maze Learning; 7 for Task 6, Working Memory Span; 28 for Task 7, Probabilistic Learning and Set Shifting; and 5 for Task 8, repeated visuospatial Maze Learning.

#### Task 1: Object Sequence Learning task paradigm

The object sequence learning task assessed how efficient subjects learn the serial temporal order of four objects. A total of five sequences were performed per session. Each of the five sequences were performed 20 times (trials), which allowed for learning the sequence during the first few trials, then to exploit the learned sequence once learning completed, and to test whether subjects form prospective plans of ordinal object locations by masking objects once a sequence was learned. The task is described in detail in (Wen et al., 2026a) and outlined with brevity here. Each sequence was composed of four objects that subjects had to touch in a predefined serial order A-B-C-D. A fifth distractor object was also shown but not relevant to the sequence. On each trial all five objects were simultaneously presented at random locations equidistant to the screen center and to each other. When subjects chose an object at the correct serial position a transient yellow halo feedback was briefly presented around the object, a positive feedback tone was played, and a green slider bar progressed towards the right. When subjects chose the incorrect object, a grey halo were briefly shown, a lower pitch sound was played, and the slider position moved back to cover one-fifth of the slider bar length. One-fifth of the slider bar length was also its starting position to allow for 4 steps towards completion. Following an error choice, subjects had to retouch, i.e. choose again, the last correct object of the sequence. When subjects correctly chose all 4 objects in the right serial order the progress bar completed and fluid reward was delivered though the sipper tube in front of the subjects. Subjects were allowed to complete the sequence within a trial by maximally committing 8 errors. When subjects committed a ninth error, the five objects were taken off the screen, the trial was aborted without reward, and the objects reappeared at random locations equidistant to the center in a new trial. After trial 6, the Object Sequence Learning task included in a subset of trials a masking condition, in which objects of future ordinal positions were masked once subjects chose the first, second, or third object of the sequence. These masking positions probed for prospective working memory planning of future locations of subjects. A prior in-depth behavioral and computational analysis suggests that five of six NHPs (83%) planned ahead around 2-2.5 positions, i.e. they chose above chance invisible, masked objects (Wen et al., 2026a).

Performance of the Object Sequence Learning task was quantified using 8 metrics (**Fig. 3E**, **S5**,**S8**). Learning speed and the number of errors committed early during learning were assessed as the (1) trials to criterion (inv.), defined as the mean trial at which a sequence was completed in three consecutive trials; (2) cumulative errors to first completion (inv.), defined as the mean number of erroneous choices made before the first completed sequence; and (3) early error count (inv.), defined as the mean number of errors during trials 1-5, when all objects were visible. Overall task performance was assessed with (4) the touch-efficiency index, calculated as the minimum number of touches required to complete a sequence divided by the number of touches made; (5) first-object accuracy, the accuracy at the first ordinal object position on Trials 6-20; and (6) pre-mask accuracy, the accuracy of visible choices made before mask onset on Trials 6-20. Sequence performance was further summarized by assessing (7) the mean maximum consecutive correct, the mean longest run of consecutive correct choices across all trials; and (8) perfect-trial rate indexing the proportion of post-learning masked trials completed without an error.

#### Task 2: Effort Control task paradigm

The effort-control task assessed how willing subjects are to exert effort for obtaining reward. Each trial of the task had two phases (**Fig. 5A**). In phase 1 subjects had to choose one of two options that varied in the amount of effort that needed to be exerted if that option was chosen, and in the amount of fluid reward subjects would earn when exerting that effort. Each option was visualized in the form of balloon outlines (5, 10, 15, or 20) indicating the number of touches needed to virtually inflate the balloon, and in the form of yellow-colored tokens (0,3,5,9,13) above the balloon outlines symbolizing the amount of fluid reward obtained when the ballon was fully inflated. Subjects had to choose among two effort-reward options presented left and right on the screen in phase 1. Phase 2 then showed only the ballon outline and token of the chosen option of phase 1 and subjects had to repeatedly touch inside the balloon outline to virtually inflate a red ballon making an air-inflating sound with each touch. The two effort-reward options varied pseudo-randomly so that low and high effort was randomly paired with low and high reward conditions, avoiding the pairing of options with identical effort and reward. Phase 2 ended with the delivery of fluid reward. Since the time to inflate a balloon dependent on the number of outlines/effort of that balloon which differed between options, the task corrected for possible differences in the duration to complete a trial for each option in phase 1 by extending the duration of the intertrial period in case the option with shorter duration (less outlines/effort) was chosen in that trial. This manipulation ensured that effort differences were linked to the number of balloon outlines and not to the time needed to complete a trial. Sessions included 80 trials of the effort control task.

Analysis modeled effort-reward choices as a function of the reward difference and the effort difference of choice options. For the first analysis of the influence of reward and the influence of effort demands on choices (**Fig. 5B**,**C**), each trial was summarized by the higher-reward option and the higher-effort option. Choices of the higher-reward option were modeled as a function of the absolute effort difference between options. Choice of the higher-effort option was modeled as a function of the absolute reward difference between options. A second analysis estimated an overall effect of VU8055 on low effort bias (high effort aversion) of choices by modeling the probability of choosing that larger-reward option as a function of the additional effort required for that reward and the reward benefit over the alternative option (**Fig. 5D**,**E**). For this analysis choices were pooled across sessions but separately for each dose condition. When additional effort was positive it indicates that the larger reward required more effort, and negative values of ‘additional effort’ indicate that the larger reward also required less effort. Thresholds were extracted at 25% and at 50% larger-reward choice probability with reward benefit held at the mean. A third analysis quantified dose effects in individual subjects on a session-by-session level by modeling session-level choices using a binomial logistic regression of the form:

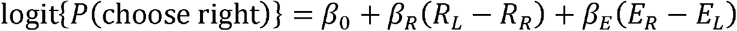

Where *R_L_-R_R_* denotes the left-minus-right reward difference and *E_R_-E_L_* denotes the right-minus-left effort difference.

The analyses provided four metrics to characterize effort control behavior: (1) ‘Reward weight’ was the metrics that indexed a subjects’ bias to choose the higher-reward option; (2) ‘effort-avoidance weight inv.’ indexing the degree of effort avoidance; the inverted sign ensured higher values reflected less avoidance, i.e. a stronger willingness to exert effort. The regression model also allowed calculating (3) the ‘reward to effort weight ratio’ indexing the relative balance of being biased towards higher reward versus lower effort choices, and (4) the ‘25% motivational breakpoint’ that indexed how much additional effort subjects accepted for making a high-reward choice before higher-reward choice probability fell to 25%.

#### Task 3: Accuracy-based Antisaccade task paradigm

We assessed inhibitory control of peripheral spatial attention and spatial interference (incongruency Simon effect) with an antisaccade task (**Fig. 2E**). The task and naming convention was adopted from human studies that found accuracy-based metrics of the antisaccade task has high consistency within subjects (Draheim et al., 2021; Burgoyne et al., 2023). Each trial began by touching the trial-start button, followed by a brief preparatory tone (0.2 s) and 0.5 s later a transiently flashed peripheral cue (an asterisk symbol, 0.15 s), followed by a briefly presented target object (0.3 s). Cue-to-target stimulus onset asynchrony was set to 0 s. All timing parameter value were established during the training period of the task prior to the experiment to elicit reliable antisaccade effects across subjects. In pro-saccade trials, the target appeared at the cued location. In anti-saccade trials, the target appeared opposite the cue, requiring subjects to avoid attentional capture by suppressing the cued-location response and identify the target at the opposite location. Following 0.5 s after target offset, the target was presented among four objects at equidistant locations to the center of the screen, and subjects had to choose the target object from this response array. In spatially congruent trials, the target in the response array appeared on the same side as the preceding target location. In incongruent trials, it appeared on the opposite side. The Antisaccade task was performed in each session 50 antisaccade and 50 pro-saccade randomly intermixed trials. Target and response object locations randomly varied across trials. Performance of the antisaccade task was scored with 12 different metrics (**Fig. 2G**,**H**, **S7B**) that quantified (1) ‘pro-saccade accuracy’ and (2) ‘anti-saccade accuracy’, as well as the analogous reaction-time (RT) metrics: (3) ‘pro-saccade RT (inv.)’ and (4) ‘anti-saccade RT (inv.)’. We also used difference metrics that quantified the antisaccade effect as (5) ‘anti minus pro accuracy contrast’ and (6) ‘anti minus pro RT contrast (inv.)’. Spatial congruency was evaluated using (7) ‘congruent accuracy’, (8) ‘congruent RT (inv.)’, (9) ‘incongruent accuracy’, and (10) ‘incongruent RT (inv.)’, as well as the difference statistics (11) ‘incongruent minus congruent accuracy contrast’ and (12) ‘incongruent minus congruent RT contrast (inv.)’.

#### Task 4: Visual Search paradigm

We assessed attention and interference control of distraction with a visual search task that varied the number of distractors and the perceptual similarity of distractors to the visual search target object (**Fig. 2A**). Subjects were presented in the first 10 trials with a single target objects, and for another 140 trials the target was embedded in a visual array with 3, 6, 9, or 12 distractors that either shared more than 2 features with the target object (high similarity of target and distractors) or that shared 1 or less feature with the target (low similarity of target and distractors, akin to a bottom up ‘popout’ target condition). Visual search performance was scored using 8 metrics (**Fig. 2D**, **S7A**): For each similarity condition, accuracy was modeled as a linear function of the number of distractors ([3, 6, 9, 12]), which yielded a slope that represents the set size effects with shallower (less negative) slopes reflecting more effective interference control. From these regression fits, we extracted: (1) the ‘Low-similarity visual-search accuracy slope’, (2) the ‘Low-similarity visual-search accuracy intercept’, (3) the ‘High-similarity visual-search accuracy slope’, and (4) the ‘High-similarity visual-search accuracy intercept’. The same analysis was performed using choice reaction times (RTs), yielding (5) the ‘Low-similarity visual-search RT slope’, (6) the ‘Low-similarity visual-search RT intercept’, (7) the ‘High-similarity visual-search RT slope’, and (8) the ‘High-similarity visual-search RT intercept’.

#### Task 5: Visual Spatial Maze Learning

The hidden maze task assessed visuo-spatial learning, cognitive control and rule following, and by repeating three mazes in Task 8 it assessed visuo-spatial memory retention (**Fig. 4A**). The task paradigm is an adaptation of the Groton Maze Learning Task (GMLT) used in humans to assess rule-guided learning and error monitoring as part of the CogState (CGS) and MATRICS (Measurement and Treatment Research to Improve Cognition in Schizophrenia) neuropsychological test batteries (Pietrzak et al., 2008; Buchanan et al., 2011). Subjects learned the path through four different mazes by choosing (touching) tiles within a 6x6 grid of tiles. Each maze was presented for five trials, allowing subjects to learn over trials by improving the speed and accuracy reaching the end tile. A maze path connected a (yellow colored) start tile with a (blue colored) end tile via 14-17 adjacent tiles and 4-6 turns.

Each maze had a unique color of the tiles and a unique background image to allow testing their retention later in the same session in task 8. Subjects followed two task rules: ‘Choose the next tile horizontally or vertically adjacent to the last correct tile’ and ‘Re-choose the last correct trial after each erroneous choice’. A correct choice of a tile transiently turned the chosen tile green and incremented the progress of the slider on top of the screen, while an erroneous choice led to a brief black-coloring of the chosen tile and the stepping back of the slider progress bar. When subjects committed multiple consecutive errors without re-choosing the last correct tile the last correct tile flashed 3 times in green to reorient the subject to the last correct tile again. Maze performance was scored with 13 performance metrics (**Fig. 4E**, **S9**): for each completion of the maze path, the error rate was calculated as the total number of erroneous tile selections divided by the number of tiles in the optimal path. Novel-maze learning was quantified using (1) ‘new first-trial error rate (inv.)’, which was the error rate during the first completion of a novel maze path; (2) ‘new last-trial error rate (inv.)’, which was the error rate on the final valid traversal; and (3) ‘new mean error rate (inv.)’, which was the mean of the trial-level error rates across all valid novel-maze traversals. Error types were further characterized using three path-length-normalized metrics calculated across the parsed maze blocks. (4) The ‘rule-abiding error rate (inv.)’ was the number of incorrect selections of an unvisited tile horizontally or vertically adjacent to the last correct tile, divided by the summed optimal path length. These choices abided to the movement rule but did not correspond to the next tile on the correct path. (5) The ‘rule-breaking error rate (inv.)’ was the number of selections classified as rule-breaking, including selections of nonadjacent tiles and selections made after an error without first returning to the last correct tile, divided by the summed optimal path length. (6) The ‘perseverative error rate (inv.)’ was the number of repeated selections of the same erroneous tile, divided by the summed optimal path length. Erroneous choices were further evaluated as (7) the ‘away error rate (inv.)’, which quantified errors in which the selected tile lay within 180° ± 90° of the vector from the last correct tile toward the end goal, indicating movement away from the goal, and (8) the ‘toward error rate (inv.)’ quantified errors in which the selected tile lay within ± 90° of this vector, indicating movement toward the goal. Each directional error rate was calculated as the number of error touches divided by the number of correct touches of the corresponding directional context. Previous studies in nonhuman primates and children have shown higher error rates when choices were away from a goal direction (Beran et al., 2015). (9) The ‘turn error rate (inv.)’ quantified errors occurring at path positions where the next correct tile required a change in direction relative to the preceding path segment. (10) The ‘straight error rate (inv.)’ quantified errors occurring at positions where the next correct tile continued in the same direction. Turn and straight error rates were calculated as the number of error touches divided by the number of correct touches of the corresponding path context. Reaction times were quantified separately as (11) ‘correct-touch RT (inv.)’ and (12) ‘error-touch RT (inv.)’, calculated by averaging the trial-level mean RTs for correct and erroneous touches, respectively. (13) ‘New duration per tile (inv.)’ was calculated as the duration of maze completion divided by the optimal path length and averaged across mazes.

#### Task 6: Working memory span paradigm

Working memory (WM) span was assessed with a task paradigm that involves continuous recognition of novel objects and the updating of a growing number of objects held in working memory with the newly chosen objects. (**Fig. 2I**). WM span was measured in 50 blocks, each taking as many trials (up to ≤15 trials) as subjects did not commit an error, i.e. each block ended with a trial in which subjects committed an error indicating their WM span for that block. The first trial of a block presented two objects. On each subsequent trial of a block, the stimulus array included all previous objects at new, random locations plus one new object. Subjects had to choose an object that they had not previously selected. Correct choices triggered a yellow halo visual feedback, a high-pitch positive feedback sound, the stepping forwards of the progress slider position on top of the screen and the delivery of fluid reward. Erroneously choosing a previously chosen object led to a grey halo around the chosen object, a lower pitch sound, the termination of the trial and block and the onset of a new trial and block with two novel objects. The task measures the number, i.e. span of objects subject can hold and update in WM and thus estimates WM capacity. Performance of the WM span task was scored with 9 metrics (**Fig. 2K**, **S7D**,**E**): (1) ‘trials to error’ indexes the average number of trials before an erroneous choice was committed. Since each trial added an object to be held in WM it indexes overall WM span performance; (2) ‘object load at 75% accuracy’ is the average number of objects held in WM at which a subject is correct in 75% of the trials; (3) ‘accuracy load slope’ is the slope of a regression of the average accuracy across trials. A shallower slope indexes a higher accuracy over later trials in which more objects are held in WM. The regression was also fit over the reaction time changes across growing number of objects, quantified in the metrics (4) ‘RT load slope’ and (5) ‘RT intercept’. Finally, overall reaction times were indexed as (8) the ‘mean RT across stimulus loads’. When subjects committed an error, they chose an object in the array that was chosen in one of the earlier trials. The erroneously chosen object could have been introduced more recently, or it could have been in WM for a longer time when it was introduced in earlier trials. Non-selective muscarinic antagonist scopolamine can affect more remote than recent WM representation (Myers et al., 2002). To assess this WM recency effect, we quantified the recency of the erroneously chosen object by calculating the average n^th^ position back at which an erroneously chosen object was introduced relative to the error trial and fitting a regression across this n-back error object distribution (**Fig. 10B**). The regression was used to extract the metric (6) ‘n-back distribution slope’ with a steeper slope reflecting a stronger WM recency effect, i.e. better WM retention for more recently updated items. The n-back slope differs when the error was committed earlier in a block and only few previous trials are available) or when errors were committed later in a block when more objects were in WM.

#### Task 7: Probabilistic Learning and Set Shifting task

We assessed flexible learning and set shifting and attention sets with a Probabilistic Learning and Set Shifting task (**Fig. 1B**). On each trial, subjects chose one among three multidimensional objects that differed in features from two dimensions, such as differing in colors and shapes. One of the objects contained a target feature value (e.g. a pyramidal shape) that was probabilistically rewarded within a block (p = 0.85), and subjects learned through trial-and-error this target feature predicting reward. A rewarded choice led to a yellow halo serving as visual feedback around the chosen object and a high-pitch tone, and the addition of token symbols to a token bar on top of the screen. Unrewarded choices led to a grey halo around the chosen object, a lower pitch tone and the removal of already attained tokens from the token bar. The rewarded target feature remained stable within a block of 27-40 trials and changed un-cued for the next block. A block started with 3 tokens and 5 empty placeholders for tokens that needed to be filled to cash out tokens for fluid reward. Blocks differed in the type of feature rule change and the reward and punishment schedules. The target feature could remain in the same feature dimension as the previous block (intra-dimensional shift, ID) or shift to another feature dimension (extra-dimensional shift, ED). Blocks could reuse the same object features (SAME, indexing set shifting) or introduce new object features (NEW, indexing new learning of an attention set). Correct choices produced either a lower-token gain (G2) or a higher-token gain (G4), whereas errors produced either a lower-token loss (L1) or a higher-token loss (L3). The experimental conditions were pseudo-randomly assigned in each session to 17 blocks of the task. Only blocks 2-16 were analyzed for the SAME/NEW and ID/ED contrast as their block transition was defined relative to the immediately preceding block.

Performance was evaluated using 28 metrics (**Fig. 1D-H**, **S6**) organized into three principal categories: learning speed, plateau accuracy, and perseverative errors. Across all eligible blocks, we calculated (1) ‘overall trials to criterion (inv.)’, defined as the first trial within a block at which mean accuracy reached at least 70% within a 10-trial forward window comprising the current trial and up to the next nine trials. Blocks in which the criterion was not reached were assigned a missing trials-to-criterion value. The sign of this metric was inverted so that higher values indicate faster learning. The metric (2) ‘overall plateau accuracy’ was calculated as the mean proportion of correct target choices from the trial following criterion attainment through the end of the block. When criterion was not reached, plateau accuracy was calculated across the entire block. We calculated (3) ‘overall CEn error proportion (inv.)’ indexed perseverative errors by counting correct responses that were immediately followed by at least two consecutive errors, divided by the number of trials within a block. This metric was also sign-inverted so that higher values indicated fewer perseverative error sequences. These three metric categories were additionally calculated separately for each of eight experimental conditions—NEW, SAME, ED, ID, G4, G2, L3, and L1—yielding 24 condition-specific metrics. Thus, separate trials-to-criterion, plateau-accuracy, and CEn-error-proportion metrics were obtained for each condition, without calculating difference or contrast scores. NEW and SAME indicated whether the relevant information changed or remained the same relative to the preceding block; ED and ID represented extra- and intra-dimensional shifts, respectively; G4 and G2 represented gains of four versus two tokens for correct choices; and L3 and L1 represented losses of three versus one previously earned tokens following errors. Finally, the 28^th^ metric ‘overall choice RT (inv.)’ quantified mean choice reaction time across trials and was sign-inverted so that higher values indicated faster responding.

#### Task 8: Visual Spatial Maze Learning with repeated mazes

To test visuo-spatial memory formation we showed in task 8 three mazes from task 5 of the same experimental session plus two new mazes. The repeated mazes had identical start and end tile positions, path, color of tiles, and background context as the same maze subjects performed earlier in the session. The two new mazes of task 8 had identical difficulty levels (number of tiles per path and number of turns) as the other mazes within the session and were used to control for the time in the session when evaluating the memory effect for repeating mazes. Task 8 provided 5 performance metrics (**Fig. 4F**, **S8**): (1) ‘repeat first-trial error rate (inv.)’, (2) ‘repeat last-trial error rate (inv.)’, and (3) ‘repeat mean error rate (inv.)’. (4) The metric ‘repeat route before first error’ quantified the proportion of consecutive correct tile-choices before the first error, averaged across repeated-maze completions. Finally, the metric (5) ‘repeat duration per tile (inv.)’ calculated maze-completion duration divided by the optimal path length and averaged across valid repeated-maze traversals.

### Statistical analysis

Analyses were performed using custom code written in MATLAB R2024b (MathWorks) and Python 3.12.6. Linear and generalized linear mixed-effects models were fitted using MATLAB’s fitlme and fitglme functions and Python’s statsmodels package.

For each task and subject, the performance metrics were extracted from each session and z-scored, resulting in one value per session, subject, dose, task, and metric before analysis using mixed-effects modeling. Overall, 85 session-level metrics were compared between vehicle and dose conditions (**Fig. S5**). Metrics where lower raw values would reflect better performance, such as reaction time, error rate, trials to criterion, or perseverative-error ratio, were sign-inverted as indicated by the abbreviation ‘inv.’ in the description of the metrics (*see* above) and figure panels. This adjustment ensured that higher, more positive values reflected improvements of behavioral performance for all metrics.

VU8055 was administered within the same week either at 0.3, 1, or 3 mg/kg on Tuesdays and Fridays, and the vehicle was administered in dedicated weeks also on Tuesdays and Fridays, and on all Mondays, Wednesdays, and Thursdays throughout the 28 weeks of the study (**Fig. S2A**). For analysis 82, 83, and 79 sessions were available with 0.3, 1, and 3 mg/kg VU8055, respectively. M/W/Th vehicle administration included 478 sessions (with Task Set A: 237; Task Set B: 241; B: 78, F: 80, J: 83, K: 78, R: 78, S: 81), and T/F vehicle administration included 66 sessions (Task Set A: 36; Task Set B: 30; 11 per subject). Control analysis quantified across all 85 performance metrics that M/W/Th and T/F vehicle sessions did not differ in performance differences scores (**Suppl. Results** ‘*Similar performance with vehicle administration on Tuesdays/Fridays and on Mondays/Wednesdays/Thursdays’,* **Fig. S13**, **S14**). Analysis compared all vehicle days (n=544 sessions, 273 in Task Group A and 271 in Task Group B), with all available dose administered days.

### Analysis of dose effects and of absolute best/worst doses

Analysis of dose effects on each performance metric used linear mixed-effects models with dose as a categorical fixed effect and subject as a random intercept:

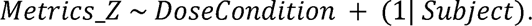

Vehicle was the reference condition, so each dose coefficient estimated the difference between that dose and vehicle. Doses were not compared with one another in these fixed-dose tests. Best/worst dose summaries used the same session-level metric values but selected the dose separately for each subject and metric. The best dose was the dose with the largest positive change relative to vehicle value. The worst dose was the dose with the largest negative change relative to vehicle. The selected best and worst values were then tested across subjects with mixed-effects models. The resulting coefficient tested whether the selected best or worst value differed from vehicle across subjects.

### Analysis of dose effects and of best/worst doses

Analysis of dose effects on each of the 85 performance metrics used linear mixed-effects models with dose as a categorical fixed effect and subject as a random intercept:

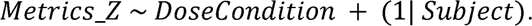

Vehicle was the reference condition, so each dose coefficient estimated the difference between that dose and vehicle. Doses were not compared with one another in these fixed-dose tests.

To evaluate whether there was a dose condition that consistently improved or impaired performance across subjects we calculated the Best- and Worst-dose using the same session-level metric values but selected the dose separately for each subject and metric. The best dose was the dose with the largest positive change relative to vehicle value. The worst dose was the dose with the largest negative change relative to vehicle. The selected best and worst dose effects values were then tested across subjects with mixed-effects models. The resulting coefficient tested whether the selected best or worst value differed from vehicle across subjects at the specific absolute dose.

We also evaluated whether there was a best dose for individual subjects regardless of whether it was the same dose across subjects. This analysis allows inferring whether individualized best doses exist for specific metrics and complement the main best-/worst-dose analysis reported in the main text. This subject-specific best dose effects analysis is shown in **Fig. S12**. For each subject and metric, we calculated the mean difference between each dose (0.3, 1, or 3 mg/kg) and that subject’s vehicle condition. The dose producing the largest improvement was selected as that subject’s best dose for the metric. We then averaged the six subject-specific best-dose effects for each metric. Two-sided one-sample t-tests assessed whether the mean effects differed from zero, and Benjamini–Hochberg fasle discovery rate (FDR) correction was applied across the 85 metrics.

### Cognitive best-/worst-dose profile

To summarize the best- and the worst-dose effects for the main metrics of each task domain we assembled a best-/worst-dose profile of cognitive and motivational effects (**Fig. 6A**). This summary used as core metrics of each task domain the high-similarity attention slopes (visual search), anti-saccade accuracy (Antisaccade), plateau accuracy and learning speed (Learning and Set Shifting), average trials to error in updating (Working Memory Span), maze learning error reduction (Maze learning), early sequence-learning errors (Object Sequence Learning), and effort-avoidance (inv.) weights (Effort Control). This analysis extracted for each metric the z-normalized best- and worst-dose effects relative to vehicle and indicates the statistical significance as obtained from the mixed-effects models.

### Analysis of the effect of subject specific baseline performance levels on dose effects

To test whether drug-induced cognitive and motivational benefits dependent on how well subjects performed the tasks at baseline, we regressed the subject specific dose effects on the baseline performance across subjects for each of 85 predefined behavioral metrics using ordinary least squares, with equal weight per subject (**Fig. S10**). Baseline was estimated from vehicle days and standardized across subjects. Sessions with dose administration were divided into alternating chronological training and evaluation subsets within each subject and dose. The best-performing dose was selected separately for each subject and metric using training sessions, and its benefit was calculated as held-out performance minus vehicle performance, scaled by the pooled within-subject standard deviation of the vehicle condition. Sessions were resampled with replacement within subject, condition, and analysis subset for 20,000 bootstrap repeats, with baseline estimation, dose selection, benefit calculation, and slope fitting repeated each time. Basic bootstrap 95% confidence intervals and two-sided tests were calculated to test for non-zero slope of the effect of the baseline performance on the dose effect strength. Bootstrap p-values were FDR corrected using the Benjamini–Hochberg procedure separately within each task. Negative slopes indicated greater benefit at lower baseline performance.

### Analysis of a task-composite dose effect

To calculate an overall net effect of each drug dose on performance across tasks, we calculated a task-composite score by averaging task-specific performance metric scores across tasks (**Fig. 6B**). We used 37 accuracy/performance-based metrics: 3 for Task 1 Object Sequence Learning, 3 for Task 2 Effort Control, 3 for Task 3 Antisaccade, 3 for Task 4 Visual Search, 7 for Tasks 5 and 8 visuospatial maze learning and repeat learning, and 18 for Task 7 Set Shifting and Learning. We did not include metrics of Task 6 Working Memory Span because it did not show task-specific effects. The task-composite scores for each session and dose were analyzed with a mixed-effects model estimating the three dose effects relative to vehicle, with subject as a random intercept. Two-sided one-sample t-tests across the six subjects tested whether the subject-level effects differed from zero.

### Analysis of subject-specific dose efficacy of VU8055 across tasks

To evaluate whether VU8055 had a net positive effect on subject’s performance irrespective of the specific tasks, we calculated subject-specific dose efficacy scores across tasks (**Fig. 6C**). These scores used the same 37 accuracy-based performance metrics used for the task composite analysis. For each subject, we compared the task-composite score at each dose with that subject’s vehicle score and then averaged the metric-level dose-minus-vehicle differences across metrics.

## Acknowledgements

We thank Nathan Traczewski and Seema Dhungana for programming support of the multitask software platform. This work was supported by the National Institute of Mental Health of the National Institutes of Health under Award Number R01MH129641 (TW). The content is solely the responsibility of the authors and does not necessarily represent the official views of the National Institutes of Health.

## Data, Materials, and Software Availability

The data and analysis code supporting this study will be available in a public repository on Zenodo upon publication. All other data are included in the article and/or supporting information.

## Author Contributions

A.N. and S.S. performed data collection; X.W. analyzed data; X.W., K.J.T, C.W.L, C.K.J, and T.W. contributed tools or analysis code for the experiment; X.W., S.S., A.N., K.J.T, C.W.L, C.K.J, and T.W. wrote and edited the paper; C.K.J and T.W. supervised the project.

