## Supplemental Information for "Broad Cognitive and Motivational Enhancement through M_4_-Selective Muscarinic Modulation"

**Affiliations:**
[1]*Department of Psychology, Vanderbilt University, Nashville, TN 37240*
[2]*Vanderbilt Brain Institute, Nashville, TN 372404*
[3]*Department of Pharmacology, Vanderbilt University, Nashville, TN 37240.*
[4]*Warren Center for Neuroscience Drug Discovery, Vanderbilt University, Nashville, TN 37232.*
[5]*Vanderbilt Institute for Therapeutic Advances, Vanderbilt University, Nashville, TN 37232.*
[6]*Department of Biomedical Engineering, Vanderbilt University, Nashville, TN 37240*

**Content**

**Supplementary Results**

- Similar VU8055 dose effects on Tuesdays and Fridays.
- Similar performance with vehicle administration on Tuesdays/Fridays and on Mondays/Wednesdays/Thursdays.

**Supplementary Discussion**

- Cholinergic Circuit Differences between Rodent and Primates.
- Working memory modulation with $M_1$ and $M_1/M_3$ receptor specific compounds and nonselective muscarinic compounds.

**Supplementary Figures S1-S18**

**Supplementary Movies S1-S7**

**Supplementary References**

### Supplementary Results

**Similar VU8055 dose effects on Tuesdays and Fridays.** The experiment alternated from one week to the next whether Task Set A and B was used on a Tuesday or Friday, which resulted in half of the Task A doses tested on a Tuesday and the other half tested on a Friday (with the same for Task Set B). We evaluated for each performance metrics how dose-effects varied by subtracting the z-scored dose effects on Fridays from the dose effects on Tuesdays.

We found there was overall a negligible effect of weekdays on the dose effects with a median absolute difference of 0.15 z values (**Fig. S13**). 81 of 85 metric estimates fell within the ±0.5-z range. Two estimates exceeded +0.5 z and two fell below −0.5 z. Dose effects metrics for Maze-learning tended to be stronger on Tuesday s(12 of 13 novel-maze metrics and all five repeated-maze metrics), although none exceeded 0.5 z in magnitude. No weekday difference survived fasle discovery rate (FDR) correction across the 85 metrics. In a separate model testing the weekday-by-dose interaction, no metric survived this correction, suggesting that dose effects did not systematically differ between Tuesday and Friday.

**Similar performance with vehicle administration on Tuesdays/Fridays and on Mondays/Wednesdays/Thursdays.** The study design administered vehicle on all Mondays/Wednesdays/Thursdays and it included dedicated weeks with vehicle administration on Tuesdays and Wednesdays that were used in other weeks as dose-testing days (**Suppl. Fig. S2**). We tested whether the vehicle testing day affected performance by comparing the 'session types' Tuesdays/Fridays and Mondays/Wednesdays/Thursdays with vehicle administration using a linear mixed effects model with subjects as random intercept and with calendar week as random intercept:

$$\text{metric\_zscore} \sim \text{session type} + (1|\text{subject}) + (1|\text{calendarweek}) ,$$

and a mixed effects model with a main effect for weekdays and subject and calendar week as random intercept:

$$\text{metric\_zscore} \sim \text{weekday} + \text{dose} + (1|\text{subject}) + (1|\text{calendarweek})).$$

We found that there was neither a statically reliable difference between these vehicle day categories when testing the session type and calendar week as random effect (**Fig. Suppl. S14**), nor was there an effect when testing for effects of weekdays (**Suppl. Fig. S15**).

### Supplementary Discussion

**Cholinergic Circuit Differences between Rodent and Primates.** Despite the functional parallels discussed in this paper between rodents and primates, there are notable anatomical differences between the cholinergic circuits across these species. In both humans and macaques, the nucleus basalis/substantia innominate complex (NB/SI) of the basal forebrain, the main source of cortical acetylcholine, is ~90% cholinergic neurons (Raghanti et al., 2011; Coppola and Disney, 2018). In contrast, cholinergic neurons make up only about 5% of the entire rat basal forebrain (Gritti et al., 2006; Coppola and Disney, 2018). Unlike the macaque cortex, the rat cortex contains a dense population (LII & III) of cholinergic interneurons (ChIs), which may compensate for the reduced basal forebrain cholinergic afferents (Houser et al., 1983; Ichikawa and Hirata, 1986; Mechawar et al., 2000).

Cross-species comparison of cognitive mechanisms is difficult due to the lack of comparative studies. In macaques, selective prefrontal acetylcholine depletion impairs working memory, but isolated NB/SI lesions do not (Aigner et al., 1991; Croxson et al., 2011). Similarly, basal forebrain cholinergic lesions in rats do not impair working memory, which has been used to suggest cholinergic depletion spares working memory (Berger-Sweeney et al., 1994; Wenk et al., 1994). However, no study in rodents specifically depleted acetylcholine in PFC to assess working memory, so this comparison is inconclusive, given rodent PFC contains its own population of ChIs.

Finally, studies in both rodents and primates suggest similar mechanisms behind sustained attention; nicotinic receptors enhance of thalamocortical input while muscarinic receptors simultaneously suppress intrinsic V1 cortical activity (London et al., 1985; Gil et al., 1997; Lavine et al., 1997; Hasselmo and McGaughy, 2004; Hasselmo and Giocomo, 2006; Disney et al., 2007; Eickhoff et al., 2007; Soma et al., 2012). There are other differences discussed by Coppola and Disney (2018), but despite species-wise differences in cholinergic innervation and sourcing, similar mechanisms may still modulate cognitively relevant circuits.

**Working memory modulation with $M_1$ and $M_1/M_3$ receptor specific compounds and nonselective muscarinic compounds.** Using a working span test, we found that the $M_4$-selective PAM VU8055 had no apparent effect on WM performance metrics in the same assay that revealed widespread improvements on attention, inhibitory control and set shifting. Consistent with the lack of a working memory effect in the WM span task there was also no modulation of the forward planning ability of subjects in the object sequence learning task when future objects were masked. Forward choices of masked objects are improved with better WM abilities in rhesus monkeys (Wen et al., 2025; Wen et al., 2026), suggesting that VU8055 did not only had no effect in the WM span task but also in the forward planning task conditions of the sequence learning task.

While the absence of an effect does not exclude the possibility that a dose outside of the tested dose range may affect working memory span, the finding suggests that working memory may more prominently be linked to other muscarinic subreceptors than the M4 mechanism. This conclusion is supported by multiple behavioral pharmacological studies in NHPs that have been robustly observed working memory deficits with systemically administered non-selective muscarinic antagonists (scopolamine, biperiden), and with agonists and positive allosteric modulators with high affinity for the $M_1$ receptor with and without affinity for the $M_3$ receptors. Among the non-selective muscarinic antagonist scopolamine impaired visuospatial paired associated performance (Taffe et al., 2002), Delayed Match-to-Sample task performance (Aigner and Mishkin, 1986;

Hironaka et al., 1992; Buccafusco et al., 2008) with delays as short as 0.5 s (Plakke et al., 2008), in DNMTS (Ogura and Aigner, 1993) and spatial delayed response tasks (Ye et al., 1999; Upright and Baxter, 2021).

Only few studies have gone beyond non-selective compound and tested how WM functions relate to muscarinic subreceptors in NHP. Most of these studies found improved WM with positive association of $M_1$, $M_1/M_3$ and possibly $M_3/M_5$ receptor activation. $M_1$ PAMs have been shown to partially recover scopolamine-induced visual spatial working memory measured (Lange et al., 2015). Behavioral pharmacology studies with agonists and partial agonists for the $M_1$ receptors (WAY-132983: $M_1$ Ki= 30 nM, phosphoinositide (PI) turnover = 88%, ED50 = 1.4 nM; $M_2$ Ki = 20 nM, cAMP = 47% of carbachol, ED50 = 600 nM) Sabcomeline), or for $M_1$ and $M_3$ receptors (compounds Cevimeline: EC50 [%EmaxAch]: $M_1$= 23 nM [82], $M_3$ = 48 nM [75]), Talsaclidine: $M_1$ = 18 nM [73], M3 = 143 nM [76]) have reported improved Delayed Match-to-Sample short term memory (O'Neill et al., 1998; Sabb et al., 1999; Bartolomeo et al., 2000; Terry et al., 2002; Heinrich et al., 2009). Antagonizing $M_3/M_5$ receptors with Descholoroclozapine impairs delayed response performance in wildtype NHPs[SS1.1] (Upright and Baxter, 2020).

**Supplementary Figures**

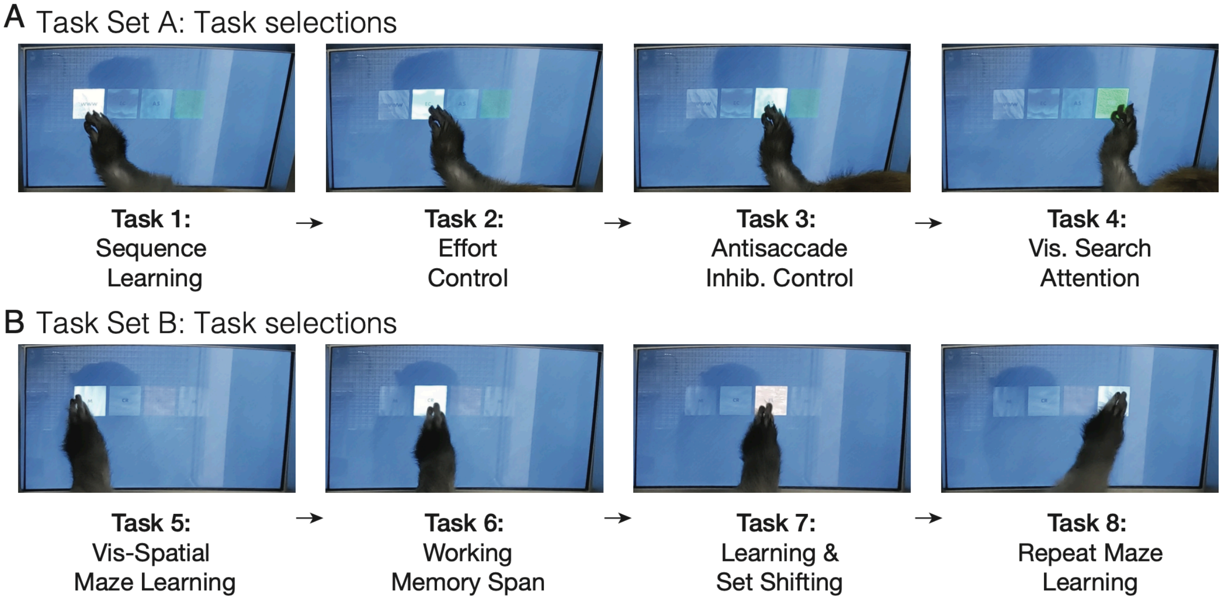

**Suppl. Fig. S1. Illustration of subjects selecting tasks for Task Set A and Task Set B.** (**A**,**B**) The photos show the task-selection (touch-) screen of the M-USE Multi-task software platform. The screens show four thumbnails for tasks 1-4 of Task Set A (*A*) and tasks 5-8 of Task Set B (*B*). The nonhuman primate sits in the apartment cage in front of the touchscreen of the Kiosk Station and selects tasks by touching the task thumbnails in a fixed order during an experimental session (from left to right). When the task selection screen is shown prior to the first task or after a task completed, only the thumbnail of the next task is fully visible while the other tasks are greyed out and inactive, ensuring subjects follow a fixed task order. Each task thumbnail has a unique structured background context theme that matches the background theme of the actual trials for that task; e.g. Visual Search is performed on a green 'grass' theme and effort control is performed on a cloud theme background (*see* also Supplementary Movies S1-7). Each thumbnail also shows the abbreviation for a task that is used in the M-USE Multi-task software platform to denote the task family of that task: 'WWW' denotes What-When-Where as the overarching task family of which the Object Sequence learning task is an instance, 'M' denotes the Maze Learning task, 'CR' refers to Continuous Recognition tasks of which the Working Memory Span task is the specific instance, and 'FL' refers to Flexible Learning tasks of which the Learning and Set Shifting task is the specific instance.

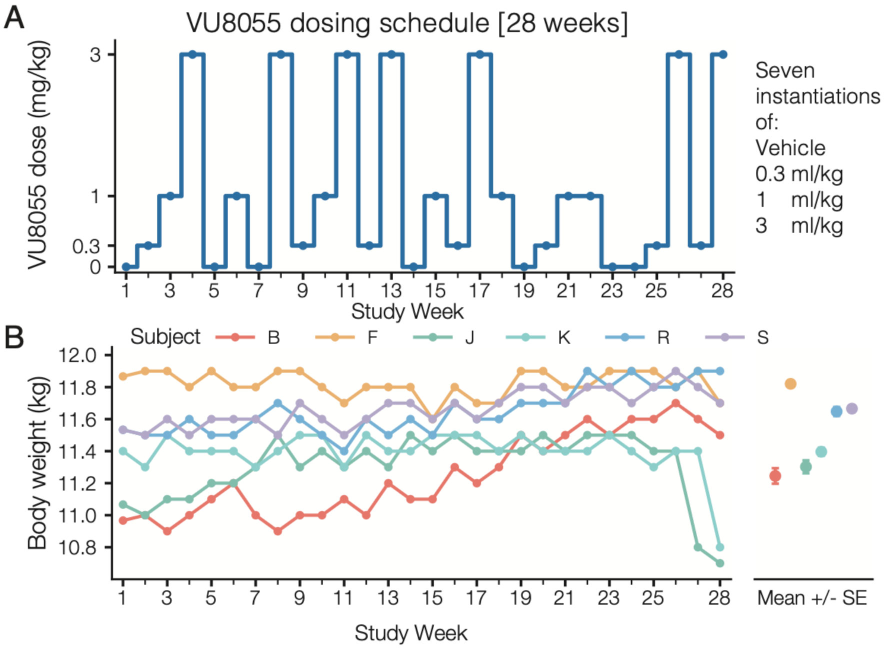

**Suppl. Fig. S2. Dose protocol and body weight stability.** (**A**) Dosing schedule of the study. In each week one dose (or vehicle, i.e. '0' mg/kg) was chosen and administered on Tuesday and Friday. Over 28 weeks the four dose conditions (vehicle, 0.3, 1, 3 mg/kg) were pseudo-randomly distributed to ensure seven instantiations per dose condition. (**B**) Body weight of each subject (colored lines) across the 27 weeks of the drug dosing study. Weights were stable and changed on average −0.01 kg (−0.1%) between the first and final dosing session. Note that the weight of two subjects (subjects J and K) dropped 0.68 kg (6.0%) and 0.27 kg (2.3%) in the last weeks after the two subjects were paused from the regular schedule due medical matters that were unrelated to the experiment. These subjects resumed study week 27 and 28 after this pause. For all other animals the 28 weeks were consecutive weeks without interruption.

NHP Adverse Drug Effect Test Battery

Effects of pharmacological challenges in NHP's will be scored on a scale of 0 or 1; where 0=no effect; normal, 1= some effect; difference noted. Baseline (pre-drug) assessments are required for determining changes within each animal. Where appropriate, a positive score will be given for increases, and a negative score will be given for decreases, in the item scored.

| Animal Name: ______________ Researcher Initials: ________<br>Session Dates: ____________ thru ____________ | | Drug Effect<br>0= No change<br>1= Some effect<br>+/- direction of change | | | | | | | | |
|---|---|---|---|---|---|---|---|---|---|---|
| Autonomic Nervous System | Definition (assessment) | Mon | | Tues | | Wed | | Thurs | | Fri |
| Salivation | Saliva on/around lips/mouth | | | | | | | | | |
| Lacrimation | Clear fluid from eyes | | | | | | | | | |
| Urination | Qualitative assessment of urine output in pan beneath chair | | | | | | | | | |
| Defecation (amount) | Qualitative assessment of fecal output in pan beneath chair | | | | | | | | | |
| Defecation (consistency) | Consistency of fecal output in pan beneath chair (hard/soft/diarrhea) | | | | | | | | | |
| Emesis | Presence of vomit | | | | | | | | | |
| Miosis | Decreased pupil diameter | | | | | | | | | |
| Mydriasis | Increased pupil diameter | | | | | | | | | |
| Ptosis | Drooping of upper eyelid | | | | | | | | | |
| Exophtalmos | Abnormal protrusion (bulging) of eyeball from the orbit | | | | | | | | | |
| Piloerection | Erection of fur | | | | | | | | | |
| Respiratory Rate | Increase/decrease in rate of inhalation/exhalation | | | | | | | | | |
| Yawn | Full extension of the jaws exposing teeth/gums | | | | | | | | | |
| Vasodilation | Dilation of blood vessels; evaluated by observing face (redness/flushing) | | | | | | | | | |
| Vasoconstriction | Constriction of blood vessels; evaluated by observing face (pale/white) | | | | | | | | | |
| Irritability | Evaluated by observation at rest and in response to mild stimulation when handled or in response to noise | | | | | | | | | |
| Body Temperature | Colonic temperature in degrees Celsius | | | | | | | | | |

**Suppl. Fig. S3. Page 1 of 2 of the Adjusted Irwin Scale used for monitoring possible side effects.** Autonomic nervous system (ANS) assessment of subjects on Mon-Fri throughout the 28 weeks of the study. Two assessments were performed daily. The first one was 5-15 minutes prior to starting the first task. The second one was immediately following completion of all tasks that day.

| | | Drug Effect<br>0= No change<br>1= Slight effect<br>+/- direction of change | | | | | | | | |
|---|---|---|---|---|---|---|---|---|---|---|
| Somatomotor Systems | Definition (assessment) | Mon | | Tues | | Wed | | Thurs | | Fri |
| Physical Appearance | Overall physical characteristics including coat consistency, skin color, affect | | | | | | | | | |
| Tremor | Involuntary movements resulting from rapid alternating contraction and relaxation of opposing muscle groups | | | | | | | | | |
| Leg Weakness | Weakness in muscle tone or resistance when extended by experimenter | | | | | | | | | |
| Catalepsy | A state of markedly diminished responsiveness in which there is a loss of voluntary motion and a plastic rigidity of the muscles; response is measured in same way as leg weakness | | | | | | | | | |
| Visuo-Motor Coordination | Determined by presenting a small treat within arm's reach of monkey and assessing ability to reach and retrieve | | | | | | | | | |
| Posture | Change in position in restraint chair; slumped/stiff/rigid | | | | | | | | | |
| Unrest | Change in activity/motor output; (e.g. restlessness, fidgeting in chair; frequent re-posturing or change in direction of movement) | | | | | | | | | |
| Stereotypies | Repeated movements; often abnormal | | | | | | | | | |
| Arousal | Degree of vigilance ranging from attentive to surrounding stimuli, hypervigilant to external stimuli or inattentive | | | | | | | | | |
| Sedation | Degree of drowsiness ranging from fully awake to slowed response to stimuli; change in posture to eyes closed or fully asleep | | | | | | | | | |
| Oral Dyskinesia | Excessive jaw movement; bruxism or tongue protrusions | | | | | | | | | |
| Bradykinesia | Stiff or slow movements ranging from normal to fixed, sustained posture | | | | | | | | | |
| Dystonia | Twisting or repetitive abnormal movement of head, neck, torso, limbs, gaping or grimacing | | | | | | | | | |
| Behaviors and Scale adapted and modified from Irwin 1968, Patel 1997 and Andersen et al 2003 | | | | | | | | | | |

**Suppl. Fig. S4. Page 2 of 2 of the Adjusted Irwin Scale used for monitoring possible side effects.** Somatomotor assessment of subjects on Mon-Fri throughout the 28 weeks of the study. Two assessments were performed daily. The first one was 5-15 minutes prior to starting the first task. The second one was immediately following completion of all tasks that day.

| Task | No. | Metric name |
|---|---|---|
| **Task 1: Object Sequence Learning** | 1 | Trials to criterion (inv.) |
| | 2 | Cumulative errors to first completion (inv.) |
| | 3 | Early error count (inv.) |
| | 4 | Touch-efficiency index |
| | 5 | First-object accuracy |
| | 6 | Pre-mask accuracy |
| | 7 | Mean maximum consecutive correct |
| | 8 | Perfect-trial rate |
| **Task 2: Effort Control** | 1 | Reward weight |
| | 2 | Effort-cost weight (inv.) |
| | 3 | Reward-to-effort weight ratio |
| | 4 | 25% motivational breakpoint |
| **Task 3: Antisaccade** | 1 | Pro-saccade accuracy |
| | 2 | Anti-saccade accuracy |
| | 3 | *Pro-saccade RT (inv.)* |
| | 4 | *Anti-saccade RT (inv.)* |
| | 5 | Anti minus pro accuracy contrast |
| | 6 | *Anti minus pro RT contrast (inv.)* |
| | 7 | Congruent accuracy |
| | 8 | *Congruent RT (inv.)* |
| | 9 | Incongruent accuracy |
| | 10 | *Incongruent RT (inv.)* |
| | 11 | Incongruent minus congruent accuracy contrast |
| | 12 | *Incongruent minus congruent RT contrast (inv.)* |
| **Task 4: Visual Search** | 1 | Low-similarity visual-search accuracy slope |
| | 2 | Low-similarity visual-search accuracy intercept |
| | 3 | High-similarity visual-search accuracy slope |
| | 4 | High-similarity visual-search accuracy intercept |
| | 5 | *Low-similarity visual-search RT slope* |
| | 6 | *Low-similarity visual-search RT intercept* |
| | 7 | *High-similarity visual-search RT slope* |
| | 8 | *High-similarity visual-search RT intercept* |
| **Task 5: Visuospatial Maze Learning** | 1 | New first-trial error rate (inv.) |
| | 2 | New last-trial error rate (inv.) |
| | 3 | New mean error rate (inv.) |
| | 4 | Rule-abiding error rate (inv.) |
| | 5 | Rule-breaking error rate (inv.) |
| | 6 | Perseverative error rate (inv.) |
| | 7 | Away error rate (inv.) |
| | 8 | Toward error rate (inv.) |
| | 9 | Turn error rate (inv.) |
| | 10 | Straight error rate (inv.) |
| | 11 | *Correct-touch RT (inv.)* |
| | 12 | *Error-touch RT (inv.)* |
| | 13 | *New duration per tile (inv.)* |

| Task | No. | Metric name |
|---|---|---|
| **Task 6: Working Memory Span** | 1 | Trials to error |
| | 2 | Object load at 75% accuracy |
| | 3 | Accuracy load slope |
| | 4 | *RT load slope* |
| | 5 | *RT intercept* |
| | 6 | N-back distribution slope |
| | 7 | *Mean RT across stimulus loads* |
| **Task 7: Probabilistic Learning and Set Shifting** | 1 | Overall trials to criterion (inv.) |
| | 2 | Overall plateau accuracy |
| | 3 | Overall CEn error proportion (inv.) |
| | 4 | NEW trials to criterion (inv.) |
| | 5 | NEW plateau accuracy |
| | 6 | NEW CEn error proportion (inv.) |
| | 7 | SAME trials to criterion (inv.) |
| | 8 | SAME plateau accuracy |
| | 9 | SAME CEn error proportion (inv.) |
| | 10 | ED trials to criterion (inv.) |
| | 11 | ED plateau accuracy |
| | 12 | ED CEn error proportion (inv.) |
| | 13 | ID trials to criterion (inv.) |
| | 14 | ID plateau accuracy |
| | 15 | ID CEn error proportion (inv.) |
| | 16 | G4 trials to criterion (inv.) |
| | 17 | G4 plateau accuracy |
| | 18 | G4 CEn error proportion (inv.) |
| | 19 | G2 trials to criterion (inv.) |
| | 20 | G2 plateau accuracy |
| | 21 | G2 CEn error proportion (inv.) |
| | 22 | L3 trials to criterion (inv.) |
| | 23 | L3 plateau accuracy |
| | 24 | L3 CEn error proportion (inv.) |
| | 25 | L1 trials to criterion (inv.) |
| | 26 | L1 plateau accuracy |
| | 27 | L1 CEn error proportion (inv.) |
| | 28 | *Overall choice RT (inv.)* |
| **Task 8: Repeated Visuospatial Maze Learning** | 1 | Repeat first-trial error rate (inv.) |
| | 2 | Repeat last-trial error rate (inv.) |
| | 3 | Repeat mean error rate (inv.) |
| | 4 | Repeat route before first error |
| | 5 | *Repeat duration per tile (inv.)* |

Overall metrics: 85
Accuracy/performance-based metrics: 67
*RT-based metrics: 18*

**Suppl. Fig. S5. Task-specific performance metrics.** The table lists the 85 session-level metrics used across the eight tasks. Metric numbering restarts within each task. Sixty-seven metrics assess accuracy or other aspects of performance, and 18 are reaction-time (RT) based; RT-based names are italicized. "Inv." indicates that the metric was sign-inverted for analysis so that higher direction-aligned values reflect better performance.

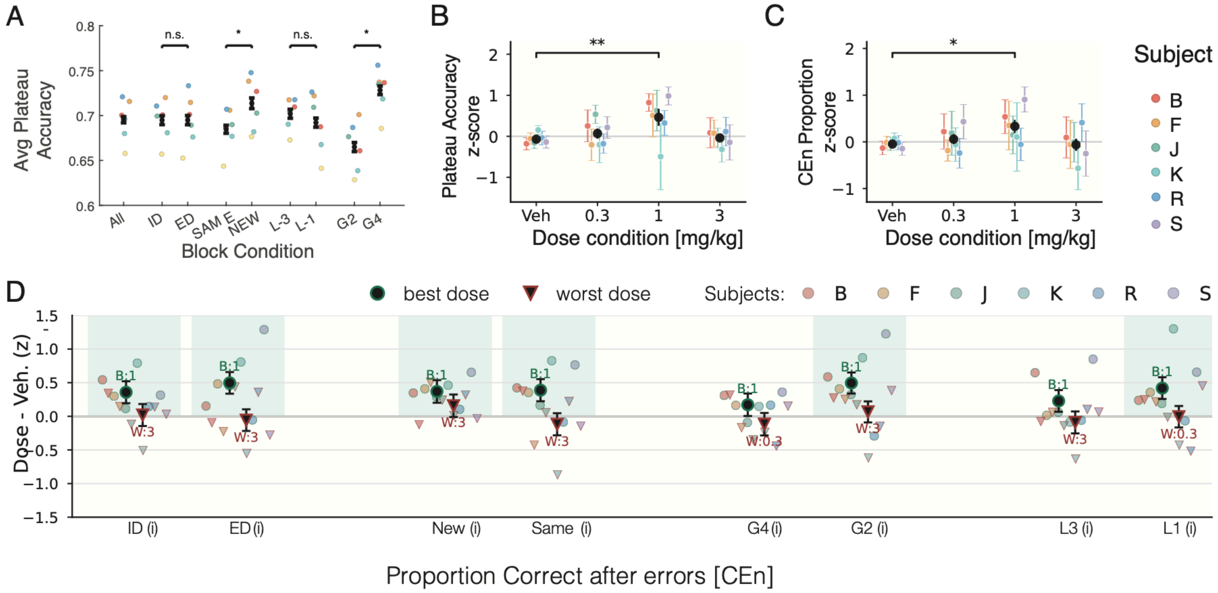

**Suppl. Fig. S6. VU8055 improved performance adjustment after errors in the Set Shifting task.** (**A**) Plateau accuracy of the set shifting and learning task was higher for NEW than SAME blocks (0.714 vs 0.685, LME of session-level avg plateau accuracy, p = 0.0064) and for higher-token gain than lower-token gain blocks (G4: 0.728 vs G2: 0.665, LME, p = 0.00020). There were no differences for block conditions ID versus ED, and L3 versus L1. (**B**,**C**) Normalized dose-response performance effects relative to vehicle baseline for plateau accuracy (*B*), and accuracy in trials following errors (*C*). Star denoted p<0.05 difference relative to vehicle based on individual selected-best-dose LME. (**D**) The norm. difference to the vehicle baseline condition and the best dose (circles) and worst dose (triangles) affecting the accuracy in trials following errors (CEn) for the block conditions intra- vs extra-dimensional shifts, new vs same object features, +4/+2 token gains, and -3/-1 token loss. Green shading denotes best doses improving CEn accuracy at p < 0.05 (individual selected-best-dose LME). There was no worst dose effect (all n.s.).

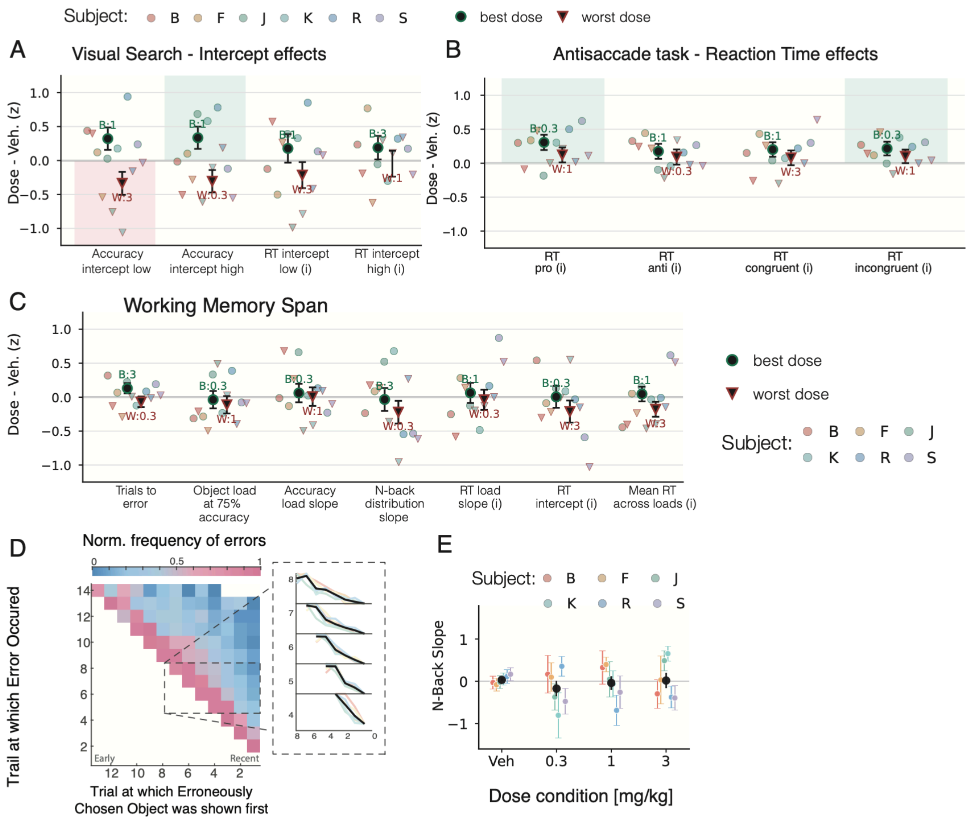

**Suppl. Fig. S7. Dose effects in Visual Search and Antisaccade task and no modulation of WM span and WM recency.** (**A**) Best dose effects on the intercept of the visual search set size effects (regression of accuracy and reaction time across increasing numbers of distractors). (**B**) Choice reaction time effects for the pro- and antisaccade condition and the spatial congruency conditions. Higher values indicate faster reaction time (**C**) Best- and worst-dose effects of the WM span task for four accuracy/performance metrics - trials to error, object load at 75% accuracy, accuracy load slope, and n-back distribution slope - and for three reaction-time metrics - mean RT across object loads, RT load slope, and RT intercept. None of these best and worse dose effects differed significantly from vehicle (all $p<0.05$). (**D**) The trial an erroneous choice was committed (y-axis) is plotted against the trial the erroneously chosen objects was introduced (x-axis). Errors were mostly based on chosen objects what were already chosen early in the block, e.g. an error on trial 8 was most likely choosing the object chosen 7 trials ago signifying confusion of an old, remotely updated object. We quantified this 'remote effect of errors' as the slope of a linear fit to the remote effect (inset on right of panel). The average of these slopes is the metric 'n-Back slope'. (**C**) Vehicle normalized dose effects on the remote error effect (*n-Back slope*) shown in B. VU8055 did not alter the recency effect (LME, 0.3 mg/kg: $p = 0.228$; 1 mg/kg: $p = 0.686$; 3 mg/kg: $p = 0.799$).

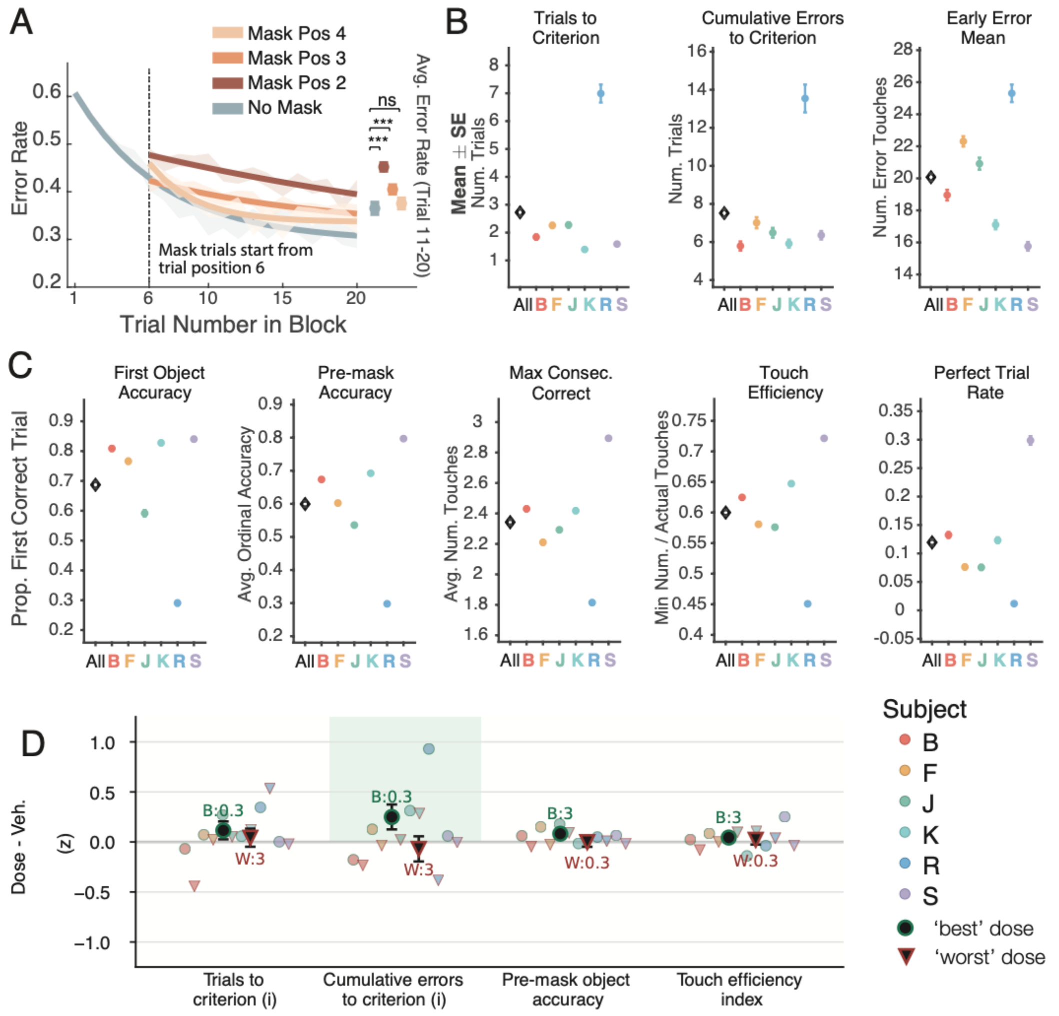

**Suppl. Fig. S8. Sequence Learning.** (**A**) Error rates declined across sequence trials without masking and when masking began before object 2, 3, or 4. At trial 20, the observed mean error rates were 30.2%, 41.4%, 35.2%, and 30.9%, respectively. The points to the right summarize trials 11–20 rather than trial-20 endpoints. (**B**) Learning measures across 274 no-drug control sessions, shown as each subject's mean ± SE and the overall session mean ± SE: trials to criterion (2.76 ± 0.14 trials), cumulative errors to first completion (7.59 ± 0.24 errors), and early error mean (20.35 ± 0.26 errors). (**C**) Corresponding performance measures: first-object accuracy (68.4 ± 1.3%), pre-mask accuracy (59.9 ± 1.0%), mean maximum consecutive correct (2.34 ± 0.02), touch efficiency (0.600 ± 0.005), and masked perfect-trial rate (12.0 ± 0.6%). (**D**) Best-/Worst- dose effect analysis. A best-dose effect was found for Cumulative Errors to Criterion (beta = 0.250 ± 0.124, p = 0.0446, session-level LME model of pooled standardized metric values, 274 vehicle and 41 low-dose sessions from 6 subjects) indexing that the low 0.3 mg/kg dose reduced erroneously choice prior to reaching the learning criterion. There were also no dose effects on performance with masks that propped prospective working memory.

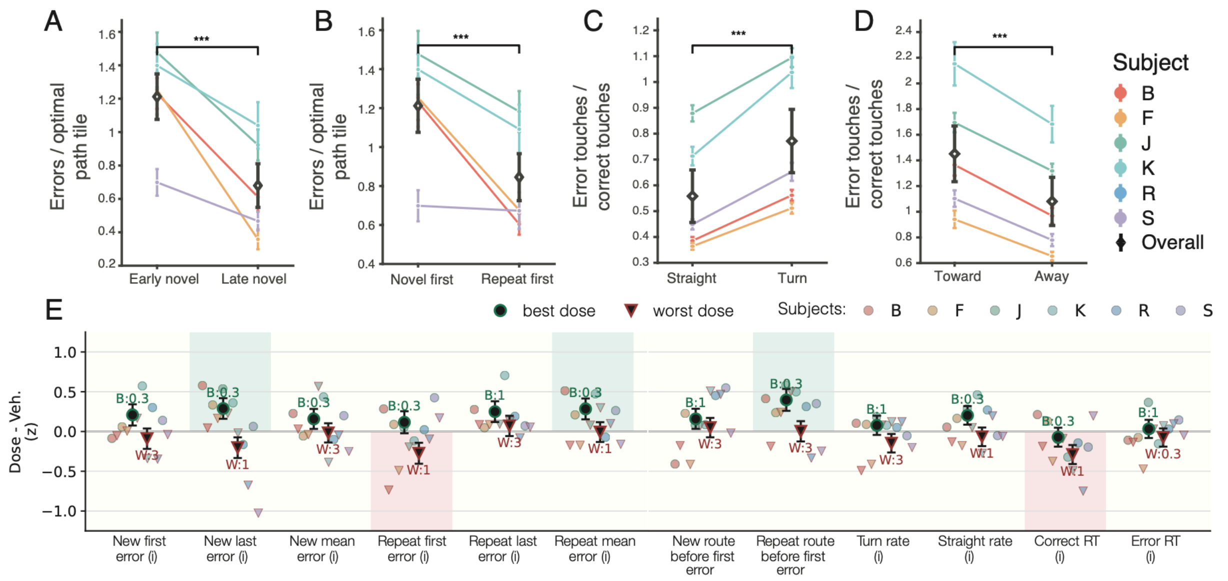

**Suppl. Fig. S9. Visuo-spatial maze learning and retention of learned routes.** Subjects learned a hidden Maze path within a block and showed first-trial savings when the Maze path was repeated in Task 8. (**A**) Early novel Maze paths (Task 5) had more errors than late novel paths (Task 8) (1.21 vs 0.68 errors per tile, p = 2.95 x 10^-14, session-level paired-difference LME model, 227 sessions, 5 subjects, same test for B,C and D). (**B**) The first path of the repeated maze had fewer errors than the same path when encountered first (first maze in repeated vs initial encounter: 0.85 vs 1.21 errors per tile, p = 2.39 x 10^-9). (**C**) Subjects made more errors at turns, where the hidden path changed direction, than on straight route segments (0.77 vs 0.56, p = 5.63 x 10^-31). (**D**) Subjects also made more errors in route segments labeled as moving toward the goal than in segments labeled as moving away from the goal (1.45 vs 1.08, p = 1.12 x 10^-14). (**E**) Best dose (circles) and worst dose (triangles) effects for metrics of the Maze learning task. Green shading denotes improvement at the best dose, while red shading denotes impairment at the worst dose (vehicle-referenced LME models with p < 0.05). At 0.3 mg/kg, subjects made fewer errors on the last traversal of the novel maze (p = 0.026) and had lower mean error rates in repeated mazes (p = 0.030). At 1 mg/kg, subjects made more errors on the first traversal of the repeated maze (p = 0.037) and showed slower correct-choice reaction times (p = 0.015). At 0.3 mg/kg, subjects also progressed farther along repeated mazes before their first error (p = 0.0039).

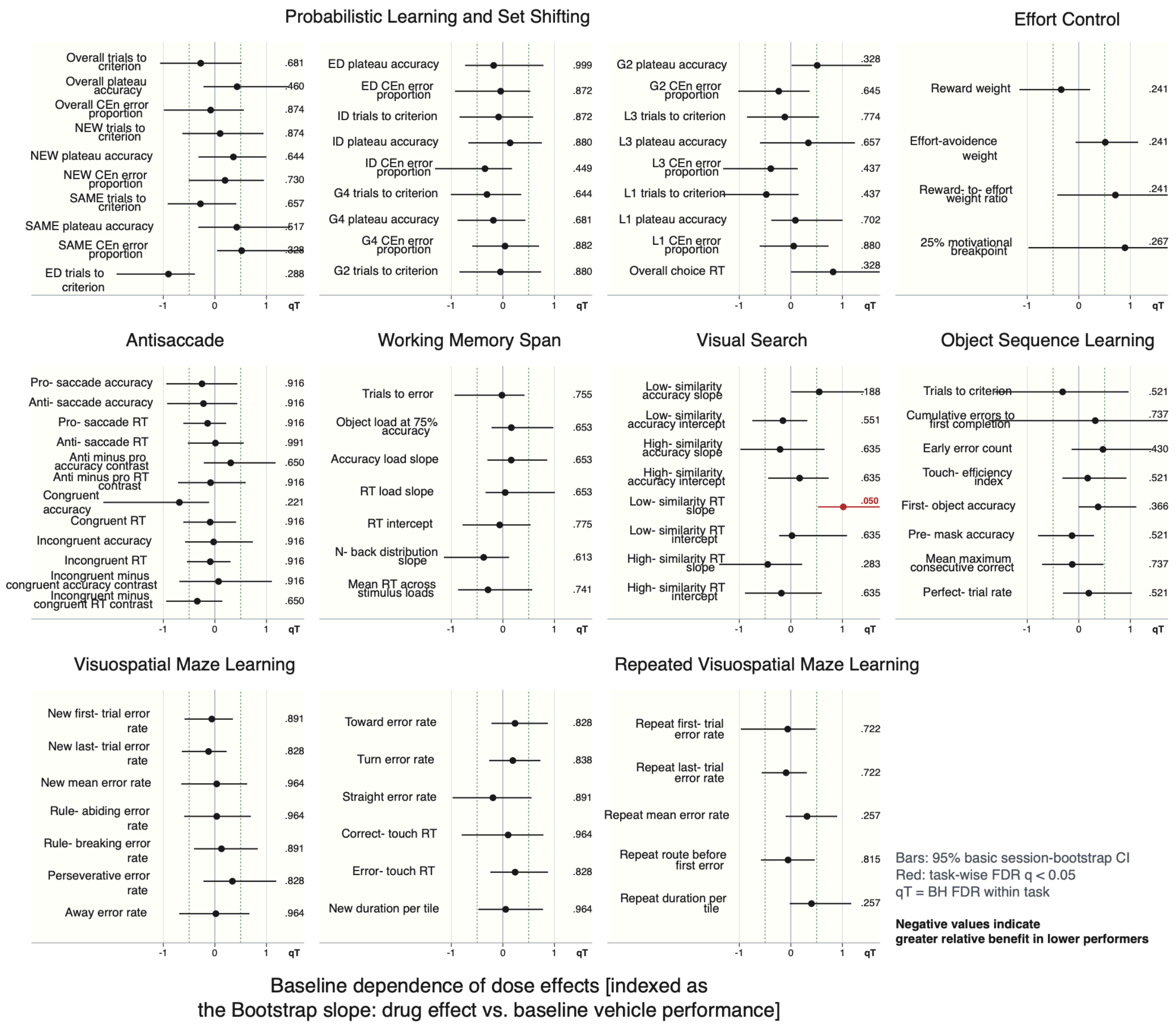

**Suppl. Fig. S10**. **No systematic influence of baseline performance on dose effects across 85 metrics.** Rows are metrics and grouped by task. Data points are average equal-weight subject-level ordinary least-squares slopes relating baseline performance to independently evaluated individualized-dose benefit (dose minus vehicle); horizontal bars are 95% basic confidence intervals from 20,000 session-bootstrap repeats. In each bootstrap repeat, sessions were resampled within subject, condition, and analysis role. Negative slopes indicate greater benefit in lower-performing subjects. The rightmost columns of each plots (qT values) show the two-sided bootstrap p-values adjusted using the Benjamini-Hochberg procedure within each task; red points indicate qT < .05. Only the visual-search low-similarity RT slope passed the task-wise FDR correction (slope = 1.017, basic 95% CI [0.536, 1.930], qT = .0496); its positive slope indicates subjects with faster baseline choice reaction times showed a stronger dose effect. No negative primary slope passed task-wise FDR.

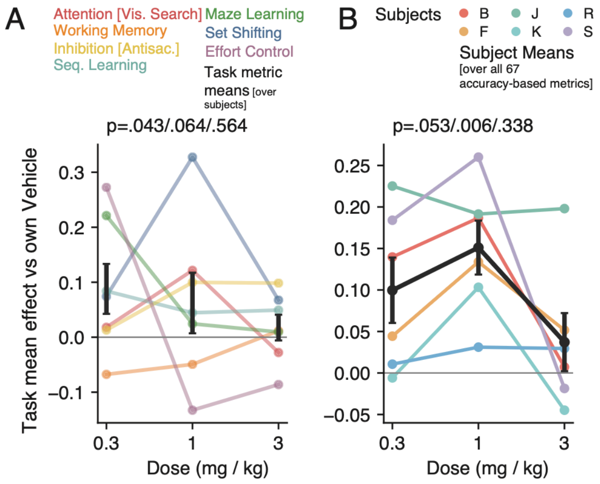

**Suppl. Fig. S11. Task-composite score and subject-specific dose effects across all 67 accuracy-based metrics.** (**A**) The task-level summary in the same format as Fig. 6B but for all 67 accuracy-based performance metrics listed in Fig. S5. (**B**) Subject-specific dose effect whosn in same format as Fig. 6C but including all 67 accuracy-based performance metrics listed in Fig. S5. Colored lines represent tasks in A and subjects in B, while black lines show the corresponding means ± SE. Displayed p values are unadjusted two-sided one-sample t-tests across six subjects, ordered by dose.

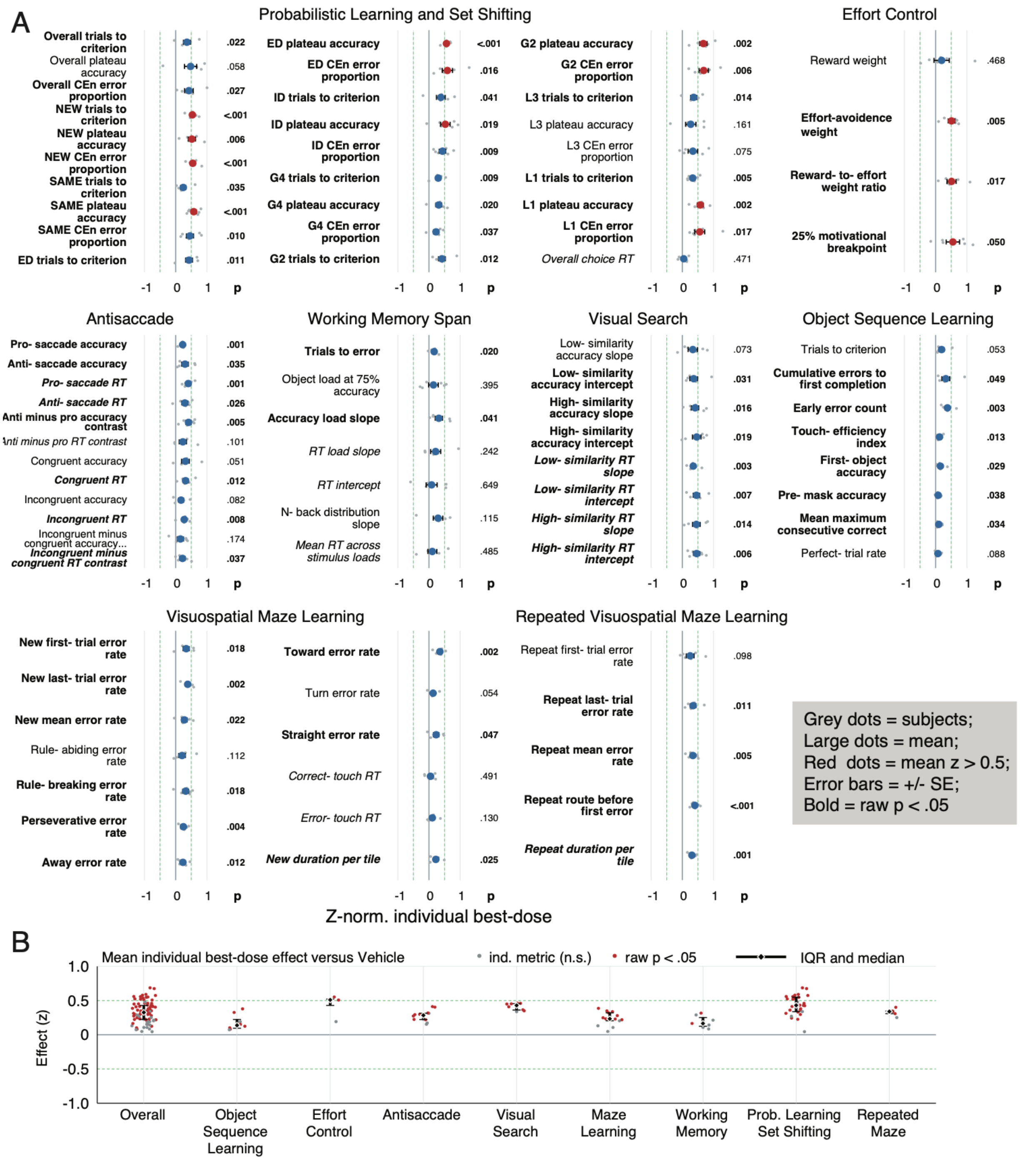

**Suppl. Fig. S12 Descriptive analysis of a subject-specific best-dose effects across 85 metrics.** (**A**) For each subject and metric, the active dose (0.3, 1, or 3 mg/kg) producing the largest improvement relative to that subject's vehicle sessions was selected independently. Small grey points show effects of the six subject; large symbols show their mean ± between-subject standard error (SE). The p values are from two-sided one-sample t-tests across subjects, and bold metric names indicate raw $p < 0.05$. (**B**) Summary of effect sizes (*y-axis*) across tasks (*x-axis*). Each point represents one metric; red points indicate raw $p < 0.05$; diamonds show the median and interquartile range across metrics. Green dashed lines mark the ±0.50 z level.

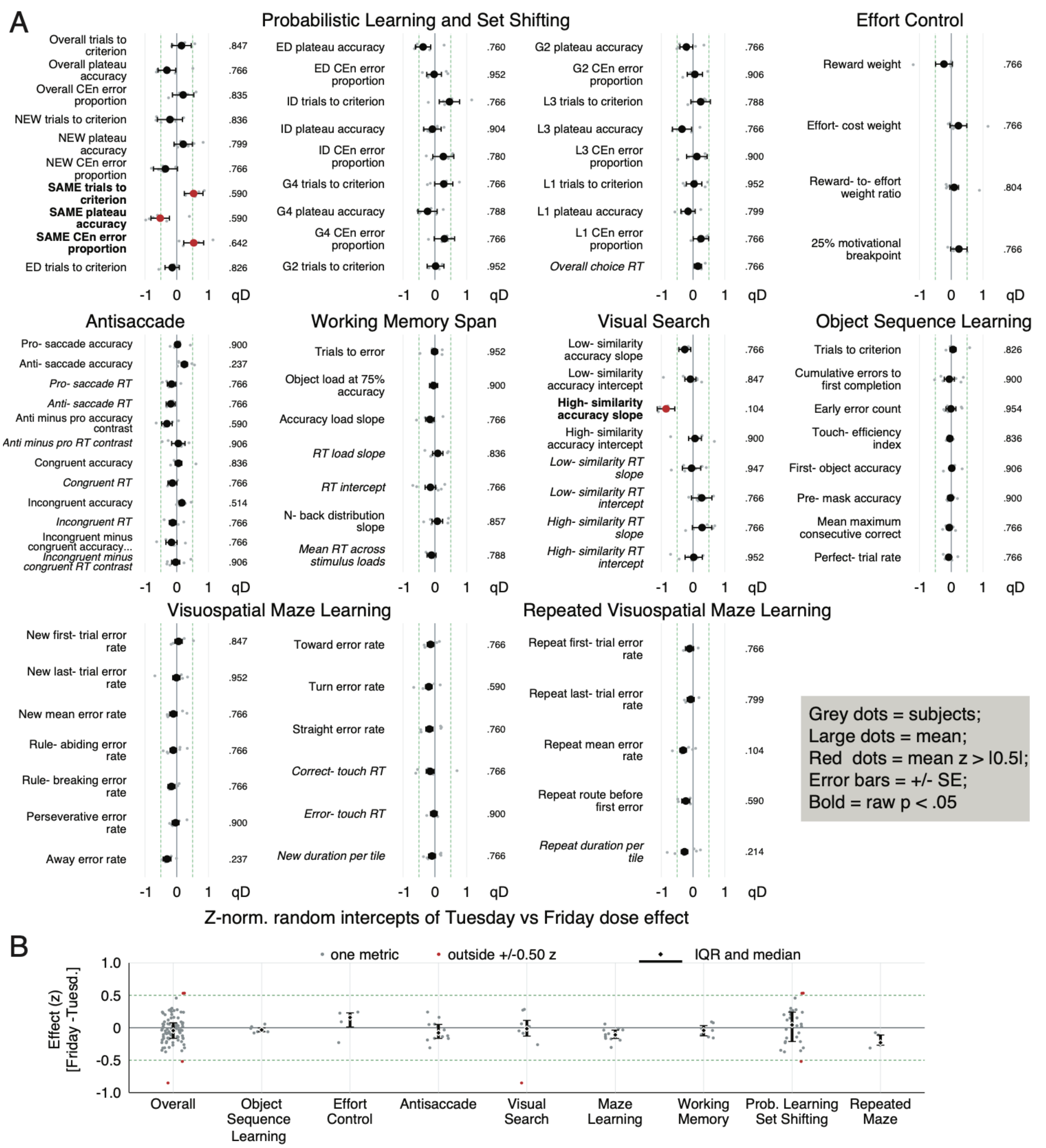

**Suppl. Fig. S13 Comparison of Tuesday and Friday dose effects.** Direction-aligned metric z scores from scheduled 0, 0.3, 1, and 3 mg/kg sessions were modeled with testing day and dose as fixed effects and crossed subject and calendar-week random intercepts. Effects are Friday minus Tuesday performance; large points and bars show model estimates ± SE, while small grey points show descriptive subject-level contrasts. The qD column gives Benjamini–Hochberg-adjusted values for the day effect across 85 metrics; dose-by-day interactions were tested separately and are not shown in this column. Red points and bold names mark estimates outside the descriptive ±0.50-z range. The lower panel shows the overall and task-wise distributions of metric effects. Neither a day effect nor a dose-by-day interaction survived correction across the 85 metrics.

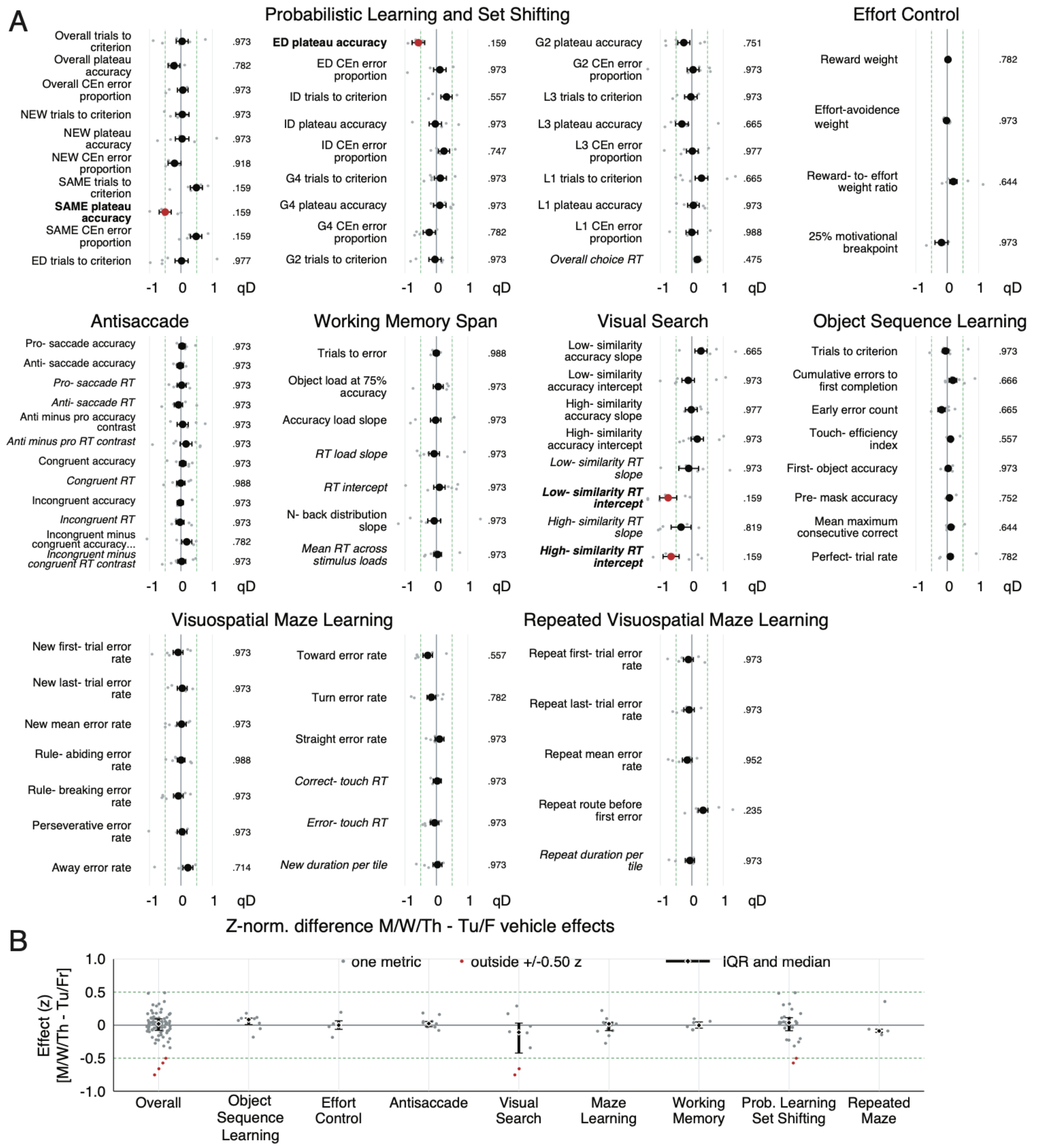

**Suppl. Fig. S15** Comparison of M/W/Th vs Tu/Fri dose effects. A and B uses the same format as Suppl. Fig. S13 and 14.

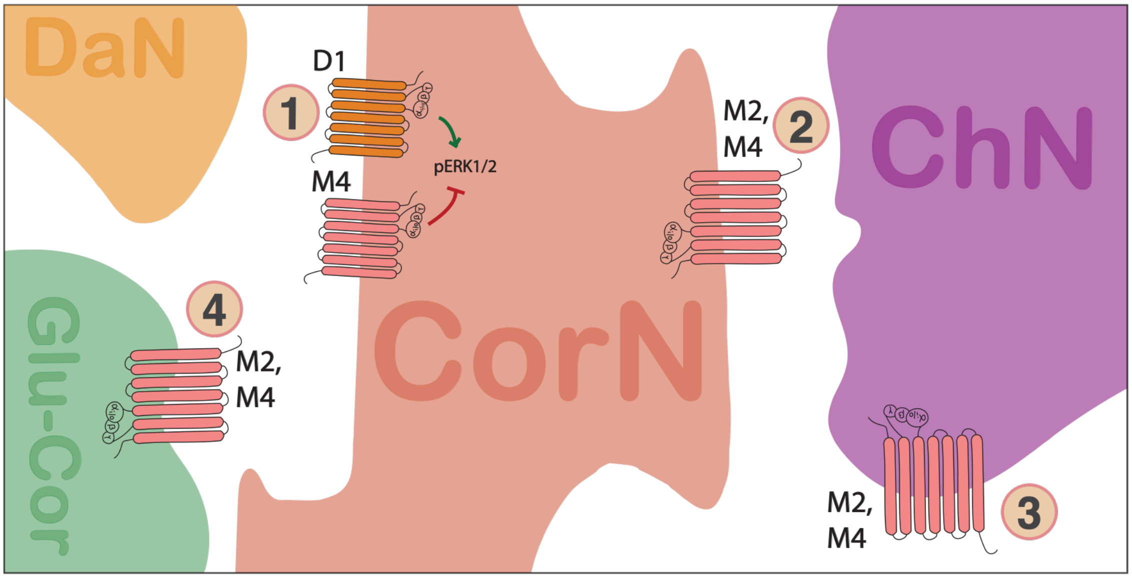

**Suppl. Fig. S16. Mechanisms of $M_4$Rs modulation in the prefrontal cortex.** Sketch of a canonical cortical neuron dendrite (CorN) synapsing to dopaminergic afferent (DaN), cholinergic afferent (ChN), glutamatergic-cortical input, and ubiquitous GABAergic input. Number point to 4 mechanisms or synaptic expression through which $M_4$ modulates cortical function across species **1:** In rats, $M_4$Rs suppress D1 mediated increase in pERK1/2 in mPFC (Xue et al., 2015; He et al., 2018). **2:** In NHPs, $M_2$Rs/$M_4$Rs are primary expressed in LIII, V/VI* (Lidow et al., 1989). They can be found postsynaptically on dendritic shafts and spines of cortical pyramidal neurons to form symmetric synapses with cholinergic axons*. **3:** $M_2$Rs/$M_4$Rs are also expressed on cholinergic axonal terminals (ChNs) to regulate acetylcholine tone in PFC and on **4:** cortico-cortico or thalamocortical terminals to regulate glutamatergic input (Glu-Cor) *(Mrzljak et al., 1993; Mrzljak et al., 1998). *Expression derived through subtracting out all receptors binding [3H] pirenzepine, which given pirenzepine IC50 for each mAchR, likely includes $M_2$, $M_3$, $M_4$, $M_5$ expression, not just $M_2$/$M_4$.

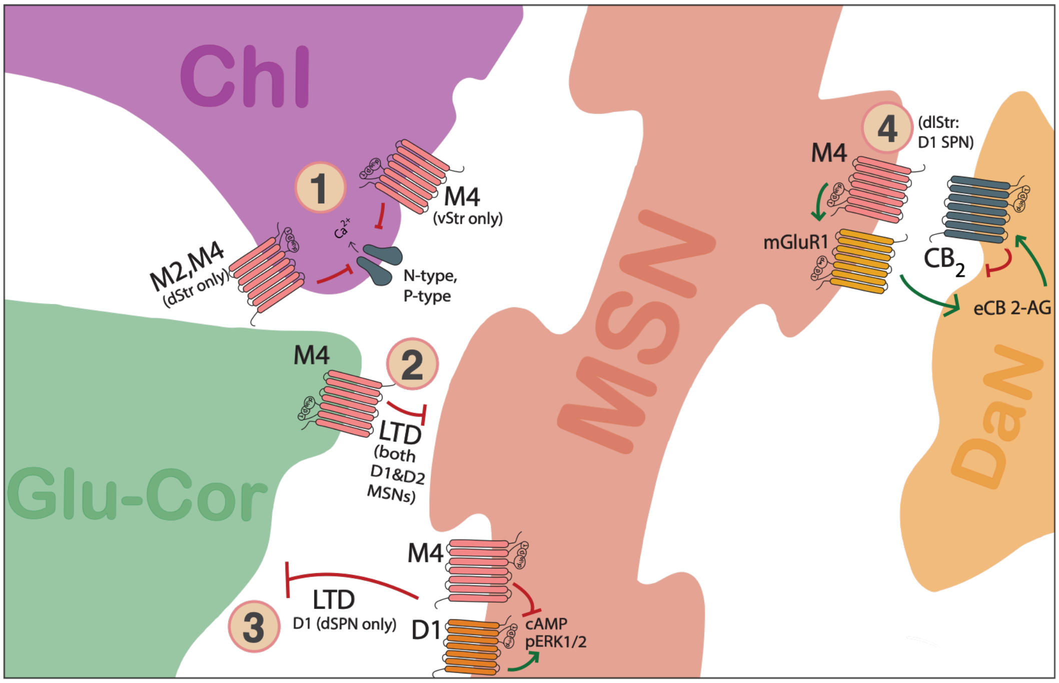

**Suppl. Fig. S17. $M_4$ regulates cholinergic, dopaminergic and glutamatergic transmission in striatum.** Sketch of the medium spiny neuron (MSN) dendrite synapsing to cholinergic interneurons (ChI), glutamatergic cortical inputs (Glu-Cor) and dopaminergic neurons (DaN). Numbers point to four mechanisms through which $M_4$ modulates striatal function: **1: (This finding is through $M_2$/$M_4$/$M_5$ knockouts or mRNA expression) $M_2$/$M_4$ on dorsal striatal and $M_4$ on ventral striatal cholinergic interneurons** regulate homo- and heterosynaptic inhibition of acetylcholine release in the striatum through inhibiting N-type and P-type $Ca^{2+}$ current (Yan and Surmeier, 1996; Threlfell et al., 2010). By regulating ACh release, $M_4$ tightly controls cortico-striatal, cortico-thalamic, and dopaminergic activity in the striatum (Tzavara et al., 2004; Threlfell et al., 2010; Downs et al., 2022). **2: $M_4$ expressed on glutamatergic cortico-striatal terminals** depress the synapse to induce long-term depression (LTD) and regulate $D_1$ and $D_2$ MSN excitability ($M_4$ mediation shown through VU152100) (Hersch et al., 1994; Ince et al., 1997; Pancani et al., 2014). **3: Postsynaptically**, in dSPNs, $M_4$ inhibits $D_1$ induced cAMP to induce long-term depression with corticostriatal synapses and attenuates $D_1$ induced ERK1/2 phosphorylation ($M_4$ mediation through VU152); the lack of this interaction enhances hyperlocomotion and behavioral sensitization following psychostimulants (Jeon et al., 2010; Shen et al., 2015; Xue et al., 2015; Ztaou et al., 2016; He et al., 2018; Nair et al., 2019). Colocalization of $D_1$ and $M_4$ is particularly high in dmStr (Garland et al., 2026). **4: $M_4$ activation of $mGluR_1$ through thalamocortical stimulation in dlStr $D_1$ SPNs** lead to sustained suppression of dopamine release through local release of eCB 2-AG in DA terminal, which activates $CB_2$ cannabinoid receptor signaling in DA terminal (VU PAM) (Foster et al., 2016; Yohn and Conn, 2018; Yohn et al., 2020). Lack of $M_4$ modulation of $D_1$ SPNs leads to increased impulsivity and increased cocaine seeking behavior, whereas lack of $M_4$ in cholinergic interneurons affected learning ($M_4$R-$D_1$RCre transgenic mice) (Klawonn et al., 2018).

| Agonists/ PAMs | Processing Speed | Attention | Working Memory | Inhibitory Control | Learning/ Set Shifting | Motivational Control | Memory |
|---|---|---|---|---|---|---|---|
| M1 | ↑⇡↺[1]○○[1]◌ | ↑⇡↺[1]○ | ↺[1]↺[1]◌ | ↺[2]↺[1]↺[1] | ↑↑↑ | | (⇡) |
| M3 | | | ↑↑↺[1]↺[1]○[1] | | ↺[3]↺[3] | | |
| M4 | ~ | ↑ | ↺[1]~ | ↑↺[1] | ↑ | ↑ | ○ |
| M1/M3 | ↑⇡○○◌◌ | ↑↑⇡⇡ | ↑⇡↺[1] | | | | |
| M1/M4 | | | | ↺[1] | ↺[1] | | |
| Nonspecific | | ↺[1]↺[1+4] | ↺[1]↺[1] | | ○ | | |

**Suppl. Fig. S18. Pro-cognitive effects of muscarinic compounds in nonhuman primates.** The table shows pro-cognitive effects of 16 muscarinic compounds with varying selectivity across $M_1$, $M_3$, $M_4$ subtypes across 7 cognitive domains reported from 19 unique studies found by searching for nonhuman primate studies testing behavior with muscarinic modulation. Symbols indicate effect in young (solid) or aged (dashed) NHPs.○ ◌ :No significant effect in either young (solid) or aged (+19) (dashed) monkeys, ↑⇡:Improvement in performance with drug in young (solid) or aged (dashed) monkeys, ↺[1]:Recovery of performance with drug following impaired performance with drug 1, (⇡):Immediate and long-term (24-48 hrs) improvement in performance in young (solid) and aged (dashed). Numbers indicate concurrent drug challenges: 1: Scopolamine, 2: Ketamine, 3: Hemicholinium-3, 4: Tacrine. *Symbols in red represent findings from this study*. Muscarinic compounds include Pilocarpine, Arecoline, WAY-132983, Compound 24, Xanomeline, VU0453595, PQCA, MK-8189, Cevimeline, Talsaclidine, L-689,660, Sabcomeline, Milameline, and RS-86.

**Supplementary Movie S1. Example performance of the Probabilistic Set Shifting and Learning task.** The movie shows example performance of multiple subjects learning which object feature is rewarded and which features are unrewarded (*see* also **Fig. 1B-D**). Its start frame depicts the touchscreen that faces the subject inside the cage-mounted Kiosk. The touchscreen displays 3 Quaddle objects and a tokenbar on top of the screen on a textured background. The tokenbar has eight placeholders for green tokens. Each block of trial starts with two green tokens occupying the tokenbar. Quaddles vary in shapes and arm styles. The movie proceeds with example trials of nonhuman primates starting trials by touching a start button, choosing an object, receiving halo feedback (grey for unrewarded, green for rewarded choices), auditory sound feedback (low pitch tone for unrewarded, higher pitch for rewarded choices), and an animation updating the token bar (subtracting available token for unrewarded and adding tokens for rewarded choices). Once tokens fill all token placeholders the tokenbar flashes white/red and fluid reward is dispensed through the sipper tube with auditory clicking sounds accompanying the turning of the reward fluid pump. The subject identity letter is indicated at the bottom of the screen.

**Supplementary Movie S2. Example performance of the Visual search task.** The movie shows performance of the initial trials showing a single target object that subjects choose to receive token reward (*see* also **Fig. 2A**,**B**). After these single-target trials, subsequent trials displays at random locations the target objects in an array of 3-12 distractors. Subjects search for and chose the target object to receive reward tokens that are saved in a tokenbar until it completes and is cashed out for fluid reward. Each trial starts by touching a start button. Objects are multidimensional Quaddle objects that share either few or ≥2 features with the target object which defines low and high target to distractor similarity conditions. The background image thema is grassland. The subject identity is indicated with a letter at the bottom of the screen.

**Supplementary Movie S3. Example performance of the Antisaccade task.** The movie depicts example performance of subjects who initiate trials, are presented with brief readiness tone, a peripheral cue and a target at the same or opposite location to the cue (pro- and antisaccade condition) and after a short delay 4 response options (*see* also **Fig. 2E**,**F**). Subjects receive token reward for choosing the target object in the display. The background image thema is water. The subject identity is indicated with a letter at the bottom of the screen.

**Supplementary Movie S4. Example performance of the Working Memory Span task.** The movie shows example performance of subjects in a block of trials that start with 2 new objects, and when subjects choose an object they haven't chosen before the subsequent trial adds one novel object also **Fig. 2I,J**). Each trial is initiated by touching a start button. The block ends when subject commit an error. The trial number at which the error is committed allows inferring the WM span. The background image thema is clouds. The subject identity is indicated with a letter at the bottom of the screen.

**Supplementary Movie S5. Example performance of the Object Sequence Learning task.** The movie shows subjects performing the Object Sequence Learning task in the different mask conditions (*see* also **Fig. 3A-C**). In one condition subjects choose objects A-B-C-D and ignore the distractor object without any masked, and in three other conditions that started from trial 6 onwards subjects are presented with the objects masks overlay objects after the first, second, or third choice during the serial choices. Correct choices were followed by a yellow halo, a feedback tone and the progress of the slider position, while were followed by a grey halo, a lower pitch feedback tone and the stepping back of the slider position. Fluid reward is delivered when the 4-object sequence is fully completed. The background image thema is rocks. Each new sequence has a different rock-style background image. The subject identity is indicated with a letter at the bottom of the screen.

**Supplementary Movie S6. Example performance of the visuo-spatial Maze Learning task.** The movie shows subjects completing an invisible path of tiles from a start to an end tile within a 6x6 grid by choosing tiles horizontally and vertically adjacent to the last correct tile (*see* also **Fig. 4A-C**). The correct tiles connecting start and end position are invisible and only briefly revealed by green color when subjects correctly choose them. This task design requires subjects to remember the green colored tile positions for subsequent trials with the same path. Subjects receive positive feedback for each choice by the stepping forward of a slider position and receive fluid reward when the slider completes. The background image thema of the maze Learning task is leather. Each maze has unique tile colors and background images. The subject identity is indicated with a letter at the bottom of the screen.

**Supplementary Movie S7. Example performance of the Effort Control task.** The movie shows subjects performing the effort control task (*see* also **Fig. 5A-E**). After starting a trial by touching a start button subjects choose either the left or right effort-reward option in Phase 1. The chosen effort-reward option then moves to the center of the screen and Phase 2 begins by subjects touching inside the balloon to virtually inflate the chosen balloon to receive the fluid reward in change for the number of tokens of the of the chosen options. The movie illustrates some trials where subjects chose the higher effort option when it is paired with more rewards, and other trials when subjects choose the option with lower effort with similar or more rewards as the alternative option. Trials of the Effort Control task uses the same sky colored background. The subject identity is indicated with a letter at the bottom of the screen.

Muscarinic Receptors in Motor Symptoms of Parkinson's Disease. J Neurosci 36:9161–9172.
